# Dynamical properties of chromatin provide insights into key folding principles

**DOI:** 10.1101/2024.05.07.592992

**Authors:** Sangram Kadam, Soudamini Sahoo, P. B. Sunil Kumar, Ranjith Padinhateeri

**Affiliations:** Department of Biosciences and Bioengineering, Indian Institute of Technology Bombay, Powai, Mumbai, 400076, India; Department of Physics and Astronomy, National Institute of Technology Rourkela, Rourkela, 769008, India; Department of Physics, Indian Institute of Technology Madras, Chennai, 600036, India; Center for Soft and Biological Matter, Indian Institute of Technology Madras, Chennai, 600036, India; Sunita Sanghi Centre of Aging and Neurodegenerative Diseases, Indian Institute of Technology Bombay, Powai, Mumbai, 400076, India; Koita Centre for Digital Health, Indian Institute of Technology Bombay, Powai, Mumbai, 400076, India

## Abstract

Even though the three-dimensional static organization of chromatin is highly studied, chromatin is a dynamic structure, and time-dependent changes are crucial for biological function. While it is known that both intra-chromatin interaction and loop extrusion are crucial to understanding chromatin organization, what is their respective role in deciding the nature of spatial and temporal organization is not clear. Simulating a model with active loop extrusion and intra-chromatin interactions, we show that under certain conditions, the measurable dynamic quantities are dominated by the loop extrusion, even though the population-averaged contact map (structure) can be dominated by intra-chromatin interactions, with loop extrusion playing no major role. Our results show that the dynamic scaling exponents with loop extrusion are consistent with the experimental observations and can be very different from those predicted by existing fractal-globule models for chromatin. We argue that one needs to measure both the structure and dynamics simultaneously to unambiguously interpret the organization of chromatin.

## I. INTRODUCTION

DNA in eukaryotic cells is packaged inside the nucleus with the help of various proteins to form chromatin [1, 2]. In the first level of packaging, DNA is wrapped around histone octamers, forming a chain of nucleosomes [2–4]. In the length-scale of genes, this chain gets further condensed into a hierarchical but dynamic structure having several loops and closely interacting topologically associating domains (TADs) [2, 5–8]. As revealed by recent experiments, chromatin organization plays a crucial role in regulating gene expression by modulating enhancer-promoter interactions [9–12]. This structural organization is thought to be a mechanism to bring multiple genes in spatial proximity (within TAD) while restricting other genes from mixing by compartmentalizing them into different TADs across TAD boundaries of varying strengths [12, 13].

The hierarchical assembly of chromatin results in interesting polymer organization and polymer properties [14]. It was proposed that the chromatin polymer can be approximated as a fractal globule, as opposed to an equilibrium globule, given its locally folded web of loops and TAD structures [15–18]. In an equilibrium globule, the overall polymer has collapsed conformation, with radius of gyration of the whole polymer *R*_*g*_ ∼ *N* ^1*/*3^, where *N* is the total number of repeat units. However, within the polymer, sub-chains of length *s < N* ^2*/*3^ can have extended configurations like a random-walk giving rise to *R*_*g*_(*s*) ∼ *s*^1*/*2^ and the scaling of contact probability as *P* (*s*) ∼ *s*^*−*3*/*2^ [16, 17]. On the other hand, the fractal globule has a non-equilibrium fractal structure with sub-chains having a similar globule structure as the overall polymer leading to *R*_*g*_(*s*) ∼ *s*^1*/*3^ and *P* (*s*) ∼ *s*^*−*1^ for all values of *s* [15–17, 19]. However, more recent studies that analyzed high-resolution data suggest that, strictly speaking, chromatin may not be a fractal globule as it shows different scaling exponents depending on the epigenetic states or different genomic size regimes and the exponents can have a distribution [20, 21].

While recent experiments have uncovered a lot about the TAD structure and other details of the static 3D organization of the chromatin polymer [2, 7, 22–25], very little is known about the dynamics of chromatin [26–30]. How this hierarchical assembly emerges and how it is maintained are questions of great current interest. The two leading hypotheses for the formation of the TAD-like structures are the active loop extrusion model and microphase separation of chromatin segments [21, 24, 31–44]. Theoretical studies have argued that the experimentally observed large chromatin loops and non-random organization are not probable by a purely thermal process and predicted an active loop extrusion process as the potential mechanism [33, 45–48]. Several studies found supporting evidence for this idea and discovered that dynamic loop formation by cohesins is crucial for chromatin organization in the interphase and CCCTC-binding factor (CTCF) plays an essential role in maintaining TAD boundaries [27, 49–57]. These findings led to the emergence of a dynamic picture where, in the interphase, structural maintenance of chromosomes (SMC) complexes like cohesins (and associated proteins) extrude a loop until they encounter a CTCF barrier. Recent microscopy experiments observed live loop extrusion *in vitro*, supporting the active loop extrusion hypothesis [51, 53, 58]. More recent imaging studies showed the dynamic nature of loop extrusion and the role of associated proteins in live cells [59, 60].

While several studies focused on the nature of extrusion process itself [61–65], a set of computational investigations have examined the role of loop extrusion in chromatin organization [21, 33, 37, 45–47, 50, 66–70]. Many of the earlier studies modeled the loop extrusion as a rate process (purely kinetic events) without explicitly accounting for the dynamics of chromatin as a polymer chain [45, 66]. They showed that the processivity of the extruders determines the loop size, coverage, and other characteristics of folded chromatin. On the other hand, polymer studies investigated the TAD formation and role of boundaries via loop extrusion [21, 33, 37, 46, 47, 50, 68, 70]. These models were used to examine the interplay between intra-chromatin interaction and loop extrusion and its role in chromatin structure and dynamics. For example, Nuebler et al. investigated loop extrusion and compartmental segregation and showed that both can affect each other in determining 3D organization[47]. Salari et al. investigated how the interactions alter the dynamics and make chromatin solid-like or liquid-like[71]. Gabriele et al. and Mach et al. studied the effect of extrusion mechanism on the dynamics of specific chromatin loci near CTCF sites using super-resolution microscopy as well as polymer simulations[59, 60]. These studies showed how dynamics at these specific sites changes in the absence of WAPL, CTCF, and cohesin.

As alluded to above, the chromatin loop extrusion has two aspects: first is the motor-like movement of cohesins, their processivity, and so on. The second aspect is the polymer nature of the chromatin itself. The chromatin polymer at the length scale of genes and TADs has a thermal relaxation time that depends on the gene/TAD size. An interplay between this polymer relaxation time and the extrusion time is likely to be crucial in determining the chromatin organization. However, there is no systematic study on how this interplay affects chromatin organization. Moreover, recent live microscopy experiments in *Drosophila* show interesting dynamic properties of chromatin (at the scale of a few genes) with non-trivial dynamical scaling exponents for relaxation time with genomic separation and mean-square displacement with time [30]. Bruckner et al. suggest that a model with a fractal globule nature and a set of dynamic rules may not explain these exponents [30]. While intra-domain attraction can form TAD-like structures, is it sufficient to get the experimentally observed dynamical scaling exponents? Is active mechanism such as loop extrusion necessary? What are the ingredients to obtain static and dynamic properties simultaneously?

In this paper, we present a polymer model for chromatin at the scale of a few TADs (few genes) and investigate if we can explain both the TAD organization as well as the dynamic exponents of the chromatin polymer simultaneously. Our model takes a specific gene region on chromosome 15 of mouse embryonic stem cells (mESC), having known CTCF boundary sites, as an example. First, we show that the model recapitulates contact map patterns, boundary probabilities, and 3D distances observed in experiments in the absence or presence of CTCF and cohesin. Then, we systematically explore how intra-chromatin interactions and various parameters of the loop extrusion mechanism (cohesin density, sliding rate, and binding/dissociation rate) affect the dynamical organization of chromatin. We show that, beyond the processivity of cohesins, the relative time scales of polymer relaxation and extrusion activity affect the chromatin organization. More specifically, we quantify how various parameters in the extrusion process affect the scaling exponents of relaxation time with genomic separation and single-locus and two-locus MSD with time. Extending our results to *Drosophila*, we show that since intra-domain attraction can dominate in determining 3D structure, the presence or absence of extrusion will be evident only when dynamic properties are measured. Our results suggest that measuring both static and dynamic properties under various conditions can discriminate the contributions from intra-domain attraction and extrusion.

## II. RESULTS AND DISCUSSION

We present results from our study of chromatin organization and dynamics resulting from the interplay between loop extrusion and intra-chromatin interactions, and argue that considering both statics and dynamics together can provide resolution to some of the puzzling aspects of 3D organization of chromatin. The model we used to simulate chromatin has the following features: chromatin is a flexible bead-spring polymer consisting of *N* beads, with each bead representing chromatin of size 2 kb (see Fig. 1a). The beads are interconnected by harmonic springs and interact with other non-bonded beads through intra-chromatin interactions such as attractive Lennard-Jones potential (see Methods section). While the self-avoiding interactions that ensure no two chromatin beads occupy the same space is inherent in our model, we present simulations with two different types of intra-chromatin interactions. One is with uniform non-bonded interaction among all beads of chromatin (all-bead uniform interaction model) representing weak non-specific attractive interactions by nucleosome tails etc. The other one is attractive interaction only among beads within a domain segment (intra-domain bead attraction model) representing specific interactions by proteins like PRC2, HP1 etc. This leads to a block copolymer model.

**FIG. 1.**
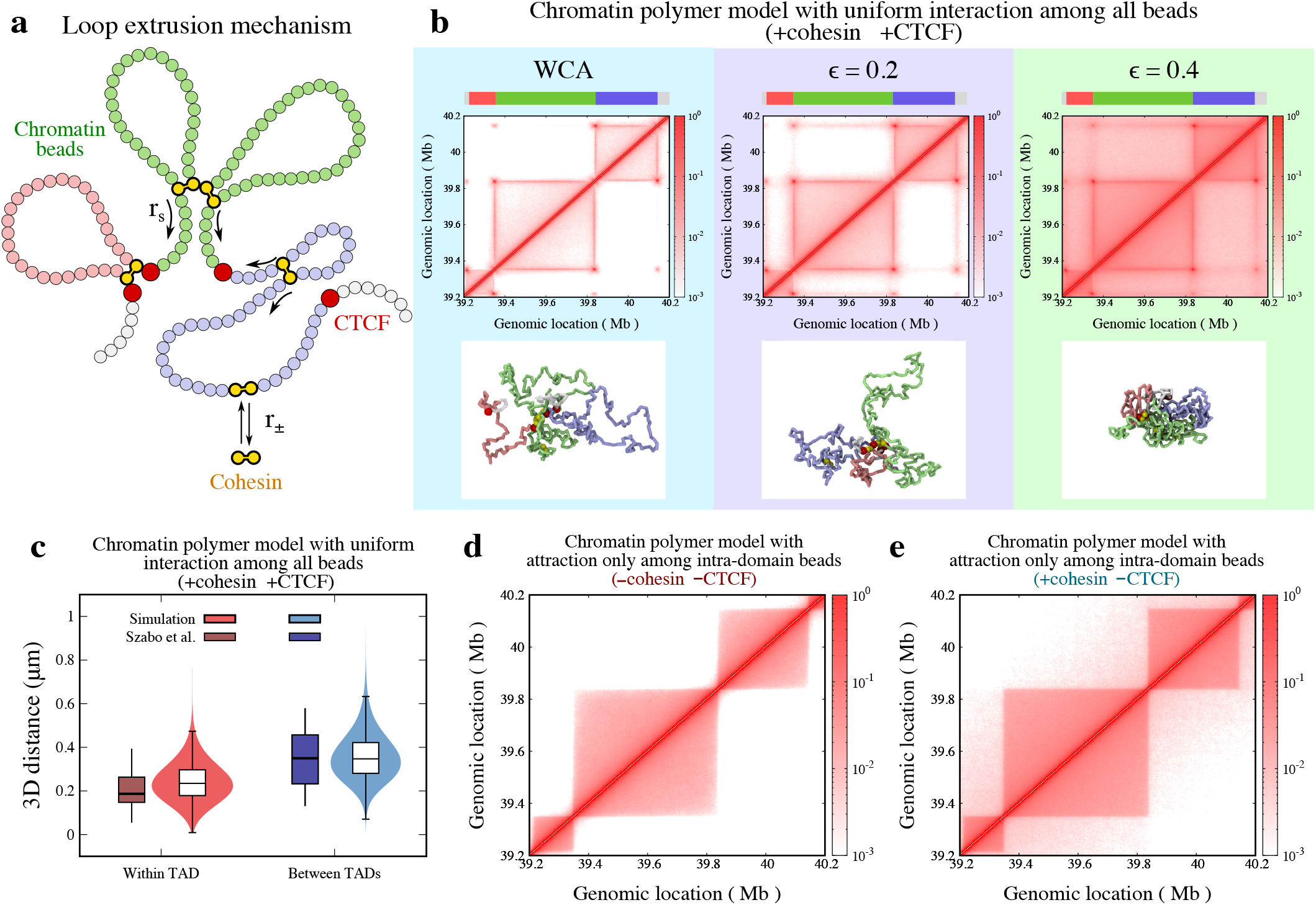
Chromatin organization with loop extrusion mechanism. **a** Schematic illustrating the loop extrusion mechanism where the cohesins (yellow color) bind/dissociate the chromatin polymer with rate *r*_*±*_. This rate determines the residence time (*T*_res_) of cohesin complexes on chromatin polymer. Two anchors of the bound cohesin slide in opposite directions with rate *r*_*s*_ on the chromatin polymer to form loops until they are blocked by the CTCF proteins (red beads) or another cohesin complex. **b** Contact maps of chromatin polymer (all-bead uniform interaction model) with loop extrusion mechanism for different intra-chromatin interactions are shown in the top row. The bottom row shows representative snapshots for respective interactions. The polymer segments are colored to distinguish different TADs, as shown in the color strip on top. The remaining parameter values are fixed 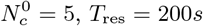 and 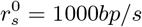. **c** The boxplot compares the 3D distance between two segments that are within the same TAD or between the neighboring TADs. The distribution of the data points from simulation (*σ* = 40nm) is plotted as a violin-plot. The 3D distance observed from our simulations is comparable to the experiments by Szabo et al. [72]. The parameter values used are 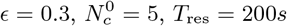 and 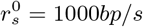. **d-e** The contact maps of chromatin polymer model with attraction between intra-domain beads only for (**d**) cohesin CTCF (no extrusion) and (**e**) +cohesin CTCF (with extrusion) conditions. The parameter values used are 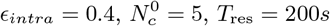 and 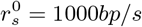.

The novelty of this work is that we are combining intra-chromatin interactions with loop extrusion, exploring the parameter space and it can lead to some interesting features as shown in the results. Loop extrusion is a fundamental mechanism involved in the dynamic reconfiguration of chromatin structure. This is governed by two sets of processes: (i) kinetic process of binding, dissociation, and energy-consuming sliding of cohesin and (ii) thermal relaxation of the chromatin polymer (see Fig. 1a and Methods). We implement kinetic rate processes such as binding and dissociation of cohesin with rate *r*_*±*_ and the sliding activity of cohesin with rate *r*_*s*_ using Monte Carlo methods [73]. During a binding event, cohesin binds to randomly chosen two adjacent empty beads. Two anchors of bound cohesins slide in opposite directions along the chromatin polymer to form a loop. This extrusion process continues until the cohesin dissociates from the chromatin polymer or encounters any barrier, such as a CTCF site or another cohesin (see Methods section). For simplicity, we assume that the CTCFs are permanently bound and can block the extrusion from both directions. This mechanism is visually represented in Fig. 1a. We vary parameters of the model like the intra-chromatin interactions (*ϵ*), the rate of extrusion (*r*_*s*_), the rate of binding and dissociation of the cohesin (*r*_*±*_), and the mean number of cohesins 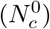 to study their effect on the chromatin organization and dynamics. We have reported all the simulation parameters in dimensionless units. Nevertheless, these dimensionless values can be converted into real units for comparison with biological systems (see Supplementary Note A). Unless specified otherwise, we choose the chromatin polymer length and CTCF locations to model a specific region of chromatin of size 1 Mb (39.2 Mb to 40.2 Mb, chromosome 15) from mESC. We consider four CTCF sites in this region at genomic locations 39.216 Mb, 39.35 Mb, 39.84 Mb, and 40.146 Mb (Fig. S1a). This region is adopted from mouse chromosome 15, experimentally studied by Szabo et al. [72].

### A. Role of loop extrusion and intra-chromatin interactions on inter-TAD and intra-TAD compaction

First, we present results from the model that considers loop extrusion and uniform intra-chromatin interactions among all beads (all-bead uniform interaction model). We plot contact maps, which visually represent the frequency of pairwise interactions between chromatin segments within the polymer. Fig. 1b shows the contact map in the presence of cohesin (cohesin+) as well as CTCF (CTCF+) for varying *ϵ*, the uniform intra-chromatin attractive interaction parameters among all beads. Here other extrusion parameters are kept constant.

The results show a consistent trend: as the attractive interaction between chromatin beads increased in strength, there was a general increase in the average number of contacts formed between any two segments (Fig. 1b). On the other hand, when non-bonded beads interact with each other through self-avoiding Weeks-Chandler-Anderson (WCA) interaction [74], there are some intra-TAD contacts and very few inter-TAD contacts. In this case, the intra-TAD contacts are primarily arising from extrusion events. The CTCF sites act as a boundary for the extrusion process and prevent inter-TAD mixing. In the case of large attractive interactions (*ϵ* = 0.4), both intra-TAD and inter-TAD contacts increase significantly. Even though there is a 1D CTCF boundary, the collapse due to intra-chromatin interactions makes 3D inter-TAD mixing possible. This shows that one-dimensional boundaries do not necessarily prevent the three-dimensional mixing of polymer segments. However, when the intra-chromatin interaction is moderate (*ϵ* ∼ 0.2), we can obtain large intra-TAD interactions and minimal inter-TAD mixing comparable to experiments (Fig. S1). To provide further clarity, we included representative snapshots of polymer conformations in the lower row of the figure. We distinguished various TAD regions using distinct colors in these snapshots, as indicated by the color strip atop the contact maps. Furthermore, red beads are used to represent CTCF sites, and cohesin complexes are illustrated in yellow. The polymer with self-avoiding WCA interaction has an extended conformation with loops and very little inter-TAD and intra-TAD mixing. The *ϵ* = 0.4 configuration points to a collapsed polymer with high inter-TAD and intra-TAD mixing. The configuration shows collapse within a TAD and very few inter-TAD contacts for the intermediate interaction strength (*ϵ* = 0.2).

We computed the 3D distances between the center of mass of segments within the same TAD and across neighboring TADs for comparison with experimental data by Szabo et al.[72] (Fig. S2a). The 3D distance between segments within a TAD is lower than the distance between segments in neighboring TADs (Fig.1c). From the simulations, we not only predict the median and interquartile ranges but also compute the full distance distribution represented as the violin plot. These results are comparable to that of Szabo et al. [72].

For the interaction parameter *ϵ* = 0.2, we investigated the role of cohesin density on the chromatin compaction. Results are depicted in Fig. S3 showing the contact map and corresponding representative snapshots for an increasing mean number of cohesin complexes 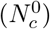. Upon close examination, it becomes evident that the overall number of contacts between chromatin segments increases with the increase in 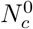, leading to higher compaction. However, this increase in contacts primarily occurs within individual TADs. Unlike the effect of intra-chromatin interactions seen in Fig. 1b, the effect of increasing the number of cohesins on inter-TAD interactions is minimal. This preservation of inter-TAD interactions is attributable to the cohesins’ predominant role in compacting individual TADs without significantly impacting the contacts between distinct TADs.

From the polymer configurations, we computed the average radius of gyration (⟨*R*_*g*_⟩), a biophysical quantity that can be measured experimentally. As expected, ⟨*R*_*g*_⟩ exhibits a decreasing trend as the strength of attractive interactions between chromatin segments is increased (Fig. S4a), suggesting stronger intra-chromatin interactions lead to a more compact polymer structure with more mixing of inter-TAD segments. Similarly, an increase in the average number of cohesin complexes also leads to the overall compaction of the polymer structure, but to a much lesser extent (Fig.S4a). Additionally, the boundary probability (*P*_*b*_) as a function of genomic location shows the strength of boundary formation at CTCF sites increases with both the strength of attractive interactions and the average number of cohesins, with the increase in 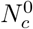 having a much stronger effect on boundary probability (Fig.S4b).

We also simulated the system in the absence of CTCF and/or cohesin. We find that, even in the presence of a constant intra-chromatin attractive LJ interaction, the TAD-like structures appear only when both cohesin and CTCF are present (See Figs. S5 and S6). Topologically associating domains (TADs) are regions (domains) where chromatin polymer is locally folded to generate high intra-chromatin interactions among the local subunits of the polymer. The boundary of these locally folded regions is typically marked by CTCF sites. These locally folded TADs have minimal interactions with nearby TADs, resulting in contact maps as shown in Fig. S5a. We observe higher inter-TAD mixing in the absence of the CTCF (Fig. S5b). However, when cohesin is absent, the contact probability values decrease drastically, implying the presence of extended polymer configurations (Fig. S5c). The TAD structure is quantified by computing the boundary probability *P*_*b*_ (Fig. S5 bottom panel). It is the probability of finding the boundary of a high contact domain at a given genomic location computed by analyzing individual polymer configurations in an ensemble (see Supplementary Note B). The case with CTCF and cohesin shows boundary peaks at CTCF sites, suggesting the presence of TADs. Next, we compared the 3D distance between segments that are within same TAD or different TADs in the absence of CTCF and/or cohesin (Fig. S7). In the absence of CTCF, we observe similar compaction with no difference in the 3D distance between segments that are within the same TAD or across two different TADs. On the other hand, the removal of cohesin leads to a significant increase in both within TAD and between TADs 3D distances, consistent with the observations from contact maps.

Unlike mammals, the evidence for domain formation through CTCF-cohesin loops is lacking in *Drosophila*[75, 76]. The domains in *Drosophila* are hypothesized to be formed through liquid-like condensate formation or micro-phase separation, where intra-domain interactions are mediated by various proteins such as PRC1/PRC2 and HP1[4, 20, 40, 77]. Although *Drosophila* lacks CTCF-dominated boundaries, it does have cohesins that can extrude chromatin loops[78– 81]. To test if loop extrusion would affect the domain structure formed through such domain-specific interactions, we simulated a chromatin model with attractive interactions only between the intra-domain beads. Fig. 1d shows that the domains can form even in the absence of CTCF and cohesin, and the presence of cohesin does not disrupt domain structure formed through the intra-domain attractions (Fig. 1e). This is consistent with the view in the field that *Drosophila* can form domains in the absence of extrusion. Such contexts are also relevant in mammals, where PRC1/PRC2/HP1-dominated intra-domain interactions compete with loop extrusion.

### B. Beyond the processivity, cohesin residence time also determines chromatin compaction

The dynamics of cohesin plays an important role in shaping the spatial structure of the chromatin polymer. Two key parameters that influence this process are the intrinsic binding and dissociation rate of cohesin 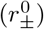 and the sliding rate 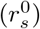. The 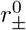 essentially controls how long a given cohesin remains attached to the chromatin polymer, i.e., residence time (*T*_res_). During the period a cohesin complex is bound, its sliding rate 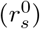 determines the amount of loop extruded. We quantify this using the processivity parameter 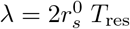, where the factor of 2 is due to the bidirectional sliding of cohesin with rate 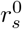. Even though some studies have investigated the role of processivity [33, 47, 66], they have not systematically explored the role of thermal relaxation of the chromatin polymer during the extrusion process. From various experiments, we know that the speed of extrusion can range from 100 bp/s to 2 kb/s [51, 53, 59, 60], while the residence time can vary from 5 minutes to 20 minutes [82–84]. Moreover, recent experiments have provided estimates for the relaxation time of chromatin segments a few hundred kb in size, ranging from 3 minutes to 15 minutes [30]. Since the relaxation time of the chromatin polymer is an additional time scale in the problem, we expect that, beyond the processivity, the relative timescales of binding/dissociation (which determines the residence time), sliding, and polymer relaxation might become important. In this section, we investigate whether the processivity solely determines chromatin organization or if the interplay between the extrusion process and polymer relaxation also plays some role. We begin by comparing the contact maps of chromatin for fixed values of processivity (*λ*), varying the residence time (*T*_res_). The range of parameter values that we explore is consistent with the experimentally observed numbers (see Supplementary Note A). In Fig. 2a, we present the contact maps for *λ* = 40kb along with different binding and dissociation rates corresponding to two residence times, which we refer as low and high residence times (*T*_res_ = 100*s* and *T*_res_ = 1000*s*, respectively). Notably, the contact maps exhibit minimal differences for fixed processivity values. The lack of full loop contacts in the contact maps (See Fig. 2a TAD corner locations) indicates that cohesins with low processivity do not reach the CTCF sites. Corresponding snapshots (see panel below the contact maps) of the polymer reveal extended conformations and the presence of smaller loops. Conversely, when we consider higher processivity (*λ* = 400kb), the contact maps show strong loop contacts emerging due to the formation of complete loops that bring CTCF sites into proximity (Fig. 2b). The associated polymer snapshots (panel below) exhibit more compacted configurations compared to the polymer with lower processivity. Although the general contact map pattern remains consistent for a fixed value of processivity, we observe a slight decrease in intra-TAD contacts for *λ* = 400kb as the residence time increases (compare TAD interiors of both the contact maps in Fig. 2b) We quantify this by plotting the average intra-TAD 3D distance (distance between beads *i* and *j* that are within the same TAD) as a function of genomic distance (*s* = | *i − j* |) between them (Fig. 2c-d). For both low and high processivity values, we observe higher intra-TAD 3D distances when the residence time is higher. This difference in intra-TAD 3D distance having the same processivity but different residence time is because of the competition between the extrusion process and the polymer relaxation. For fixed processivity, when residence time is longer, the extruded polymer will have more time to relax. Conversely, when residence time is low, the cohesin extruders dissociate and bind back quickly before the polymer relaxes, leading to higher compaction (see supplementary Movie S1). To quantify this further, we calculate the average radius of gyration(⟨*R*_*g*_⟩) of the chromatin polymer as a function of processivity (*λ*) (see Fig. 2e). ⟨ *R*_*g*_⟩ is consistently higher for *T*_res_ = 1000s (red curve), implying that longer residence times may allow the polymer to relax more. As we decrease the residence time, we observe a reduction in ⟨*R*_*g*_⟩ for all values of *λ*. The relaxation time of the polymer segment of a few 100 kb size is of the order of 1000s (see the following section for details). For a fixed processivity value, if the residence time *T*_res_ = 1000s of the cohesin is comparable to this relaxation time, the chromatin polymer has a more open structure (Fig. 2e right panel). On the other hand, the polymer forms compact domains when the residence time of the cohesin is *T*_res_ = 100s, which is much less than the relaxation time of the polymer segment (Fig. 2e right panel). Therefore, in addition to the processivity of the cohesin complex, the relative value of the residence time as compared with the polymer relaxation time also contributes to the chromatin topology.

**FIG. 2.**
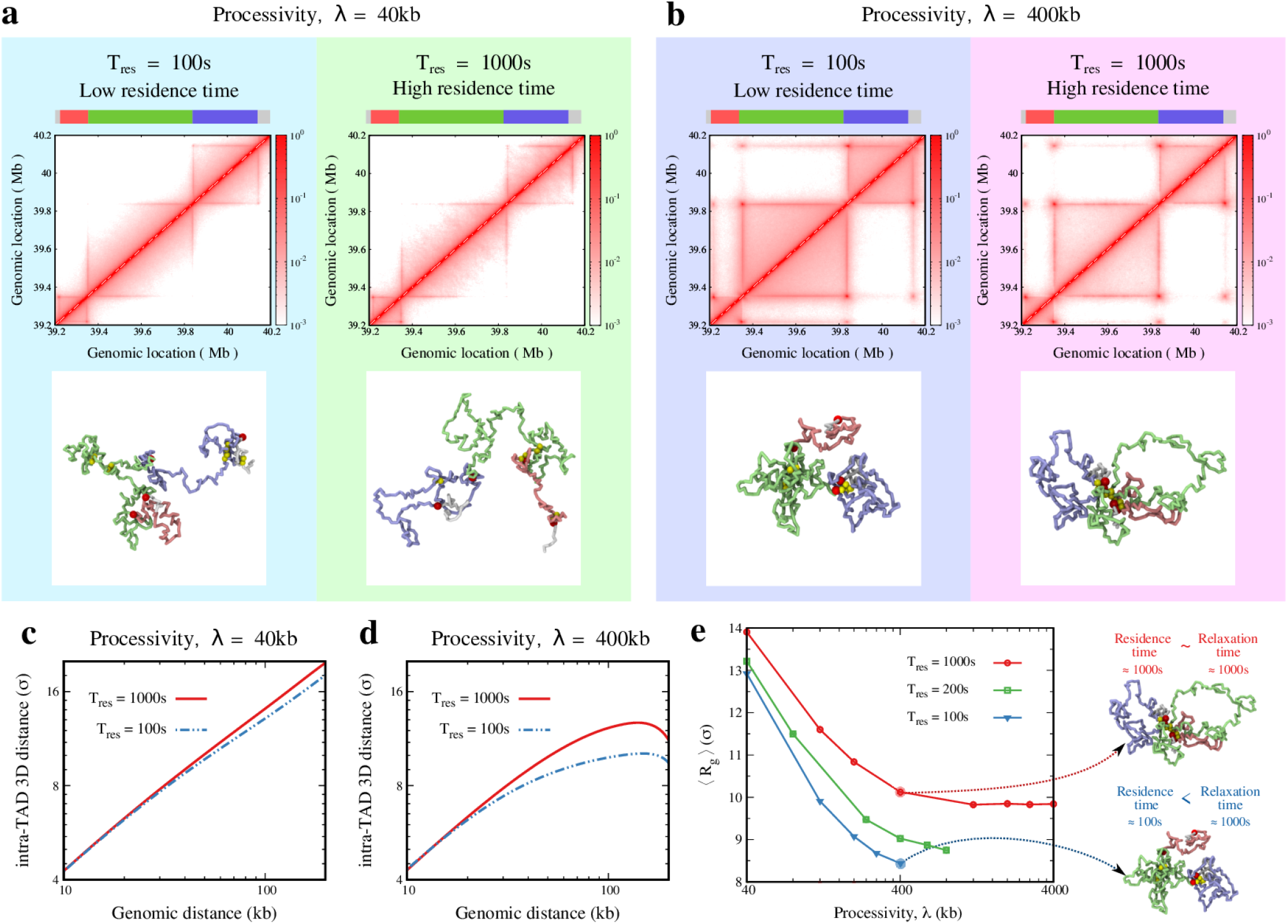
The interplay between residence time and the processivity of cohesin complexes. **a**,**b** Contact maps for different values of residence time *T*_res_ are shown for fixed processivity values (**a**) *λ* = 40*kb* and (**b**) *λ* = 400*kb*. The corresponding representative snapshots are shown below the contact maps. The polymer segments are colored to distinguish different TADs, as shown in the color strip on top. **c, d** Average intra-TAD 3D distance versus genomic distance (*s*) is plotted for different values of *T*_res_ for fixed processivity values (**c**) *λ* = 40kb and (**d**) *λ* = 400kb. **e** Average radius of gyration as a function of *λ* is plotted for different values of *T*_res_. Representative snapshots of the polymer for *λ* = 400kb are shown on the right side for *T*_res_ = 100s and *T*_res_ = 1000s. All other parameter values for all plots are fixed to intermediate values: *ϵ* = 0.2 and 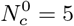.

We also compute the 3D distance between two chromatin segments within the same TAD or two neighboring TADs (see a schematic description in Fig. S2a). We choose these segments for comparison with the super-resolution microscopy experiments that measure the 3D distance between the same regions [72]. A trend similar to what is seen in Fig. 2e is observed when we examine the 3D distance between the center of mass of these segments (Fig. S2b). In this case, both the 3D distances within the TAD and between TADs are consistently higher for the polymer with longer residence time of the cohesin for a given processivity (green solid and dotted curves). These 3D distances decrease as the residence time is reduced (red and blue curves). Moreover, the 3D distance between TADs is considerably higher than the 3D distance within TAD, consistent with the experimental observations [72].

### C. Loop extrusion mechanism and intra-chromatin interactions influence the relaxation time of chromatin segments

Apart from the static spatial organization, we also quantify the effect of the intra-chromatin interactions and loop extrusion mechanism on the chromatin dynamics. First, we compute average auto-correlation function of the 3D distance vectors **R**(*s, t*) between any bead pairs *i* and *j* separated by a genomic distance *s* = |*i − j*|*σ* (see Fig. 3a) as

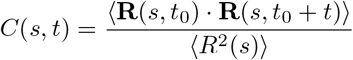

Here ⟨*R*^2^(*s*)⟩ is the mean square distance between chromatin loci separated by genomic distance *s*. The relaxation time *τ*_*r*_ is defined as the time it takes for this correlation to decay to the value of *C*(*s, t* = 0)*/e*. We perform this analysis for different values of genomic separation (*s*) and present the data for different parameter sets (Fig. 3).

**FIG. 3.**
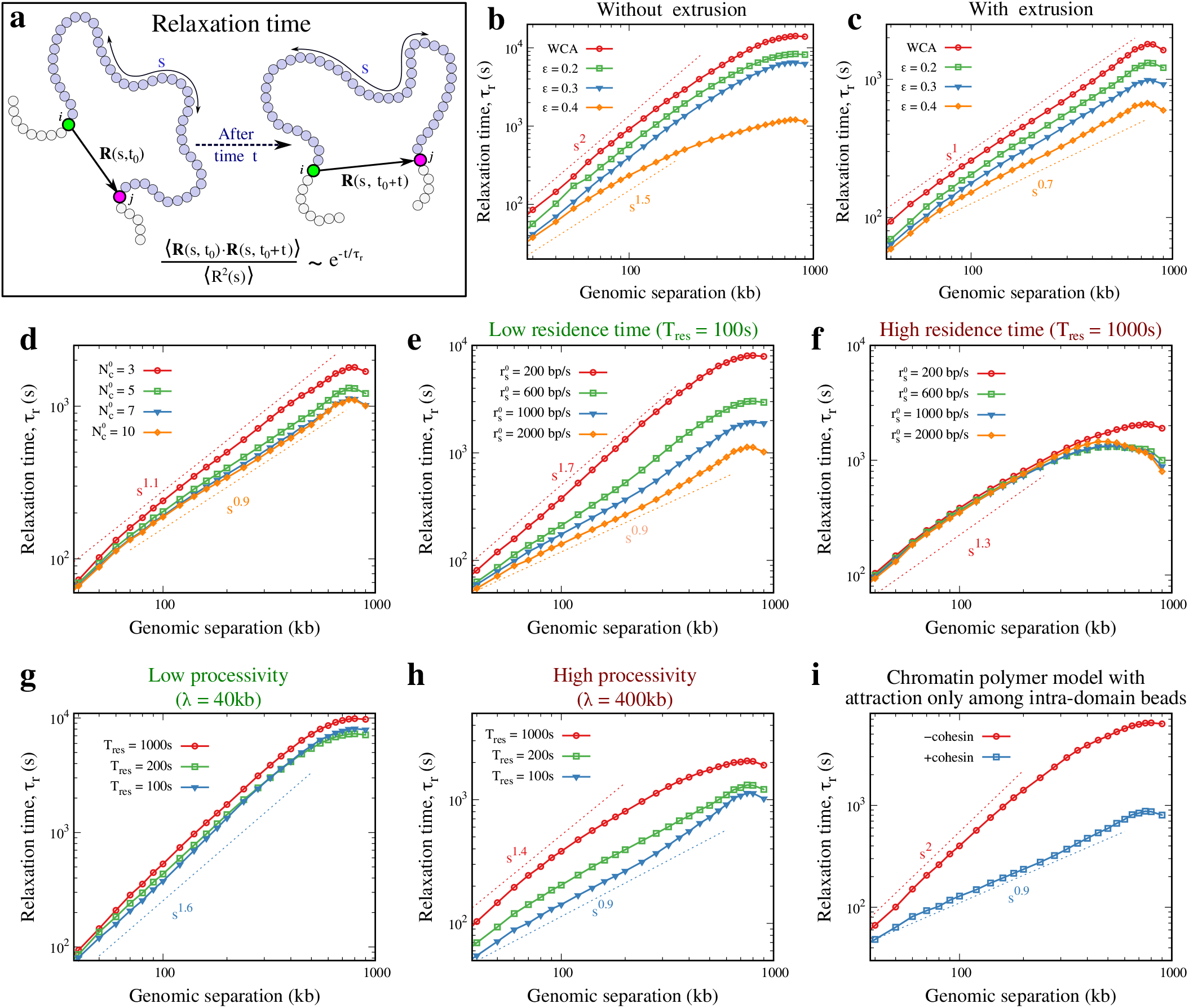
**a** Schematic showing the calculation of relaxation time for a polymer segment of size *s*. The two-locus auto-correlation function, for any two bead pairs *i* and *j* with genomic separation *s*, decays exponentially with time. **b** The relaxation time for a polymer without an active loop extrusion mechanism is plotted as a function of genomic separation *s* for different intra-chromatin interactions. **c** The scaling of relaxation time with genomic separation for a polymer with the loop extrusion mechanism is shown for different intra-chromatin interactions. **d** The plot shows how scaling of relaxation time varies with the mean number of cohesin complexes in 1Mb region. **e-f** The scaling of relaxation time with genomic separation is plotted for different sliding rates of cohesin 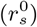 for (**e**) a lower value of residence time and for (**f**) a higher value of residence time. **g-h** The Effect of residence time on the scaling of relaxation time is compared for polymers with (**g**) low processivity and (**h**) high processivity. In all the plots (**b-h**), apart from the parameters that we vary in each plot, all other parameter values are fixed to an intermediate value: 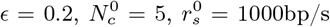, and *T*_res_ = 200s. **i** The relaxation time versus genomic separation is plotted for chromatin polymer model with attraction only among intra-domain beads in the presence and absence of cohesin. Here, the intra-domain attractive interactions is *ϵ*_intra_ = 0.4 and the extrusion parameters for +cohesin condition are 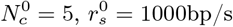, and *T*_res_ = 200s.

Theories of polymer dynamics expect a power law scaling of *τ*_*r*_ ∼ *s*^*γ*^ [85]. The Rouse dynamics predicts *γ* = 2.2 for a self-avoiding walk (SAW) polymer and *γ* = 5*/*3 for the fractal globule [85]. However, a recent experiment found that for chromatin *γ* ≈ 0.7, which is far from what is predicted for the fractal globule [30]. How to explain this surprising experimental finding is an open question. In this section, we examine how active loop extrusion and intra-chromatin interactions affect the value of *γ*.

We computed *τ*_*r*_(*s*) for our chromatin model with uniform interaction among all beads. The results are presented in Fig. 3b-h for different parameters associated with the loop extrusion mechanism. First, we consider the control case of the chromatin polymer without any loop extrusion (Fig. 3b). When attractive intra-chromatin interactions are absent (WCA), the *γ* value is close to 2. With increasing strength of attractive interactions (*ϵ* ≥ 0.2), the *γ* decreases further. Then, we investigate the role of extrusion by studying chromatin polymer by varying the extrusion parameters. In the presence of extrusion, *γ* decreases from its value in the absence of extrusion for all corresponding values of *ϵ* (compare Fig. 3b and Fig. 3c). Note that the dependence of relaxation time (*τ*_*r*_) on genomic separation (*s*) with extrusion and intra-chromatin interactions is close to what is experimentally observed [30].

We further systematically investigate how extrusion parameters influence *γ*. Fig. 3d shows the impact of varying the mean number of cohesins on the scaling of the relaxation time. We find that *γ* decreases slightly as the cohesin density increases. Next, in Fig. 3e-f, we explore the effects of altering the sliding rate and residence time of the cohesin on the scaling of relaxation time (*τ*_*r*_(*s*)). When residence time is shorter (*T*_res_ = 100s), the relaxation time decreases with increase in sliding rate, which helps the polymer reorganize faster (see Fig. 3e). Interestingly, at this low residence time, as we increase the sliding rate *r*_*s*_, the *γ* value decreases from that predicted by the fractal globule model (*γ* = 5*/*3) towards the experimentally observed value *γ* = 0.7 [30]. However, as we increase the residence time, we observe that the scaling of relaxation time appears to be independent of the sliding rate (Fig. 3f). At such values of residence times, the cohesins would form complete loops and stay at the CTCF sites, reducing the effect of the sliding rate. Similarly, we also check the impact of processivity on the value of *γ* (Fig. 3g-h). For lower processivity, residence time has minimal effect on the exponent *γ*. However, for higher processivity, the effect of residence time is more pronounced with increasing residence time, leading to an increase in *γ* and vice versa.

It is puzzling that the *γ* = 0.7 for the relaxation time is observed in *Drosophila*, where it is argued that loop extrusion is not prominent [75, 76]. While intra-chromatin interactions are considered a major contributor to domain formation in the fly[4, 77, 86], our findings reveal nuanced insights into chromatin dynamics. When intra-domain attractive interactions dominate, as shown in Fig. 1d-e, extrusion has minimal effect on the domain structure and contact map. While cohesin might play a negligible role in 3D domain structure, we hypothesize that it could significantly impact chromatin’s dynamic properties. Our analysis of relaxation dynamics of chromatin polymer in a *Drosophila*-like system shows a significant change in the relaxation time exponent in the presence of cohesin (Fig. 3i) even though the 3D domain structure (contact map, Fig. 1d-e) remains unchanged. This suggests that active loop extrusion mechanisms substantially contribute to chromatin dynamics, even in *Drosophila*. Our prediction can be tested by experimentally studying the dynamics in the presence and absence of cohesin in *Drosophila*. It should also be noted that recent experiments do report that extrusion by cohesin in conjunction with RNAPII and other proteins can form loops[79–81].

### D. Single-locus and two-locus mean square displacement reveals anomalous diffusion in chromatin

To quantify the dynamics further we compute the mean squared displacement of any bead *i* as a function of time *M*_1_(*t*) = ⟨(**R**_*i*_(*t*_0_ + *t*) *−* **R**_*i*_(*t*_0_))^2^⟩, averaged over all beads (Fig. 4a). According to the Rouse dynamics, one expects the single-locus MSD *M*_1_ ∼ *t*^*α*^, where *α* = 0.5 for ideal polymer [87]. Studies of the fractal globule polymer have suggested *α* ≈ 0.4 [88]. However, recent chromatin experiments found *α* to be varying around 0.5. In this section, we investigate how the time dependence of MSD is affected by loop extrusion and intra-chromatin interactions.

**FIG. 4.**
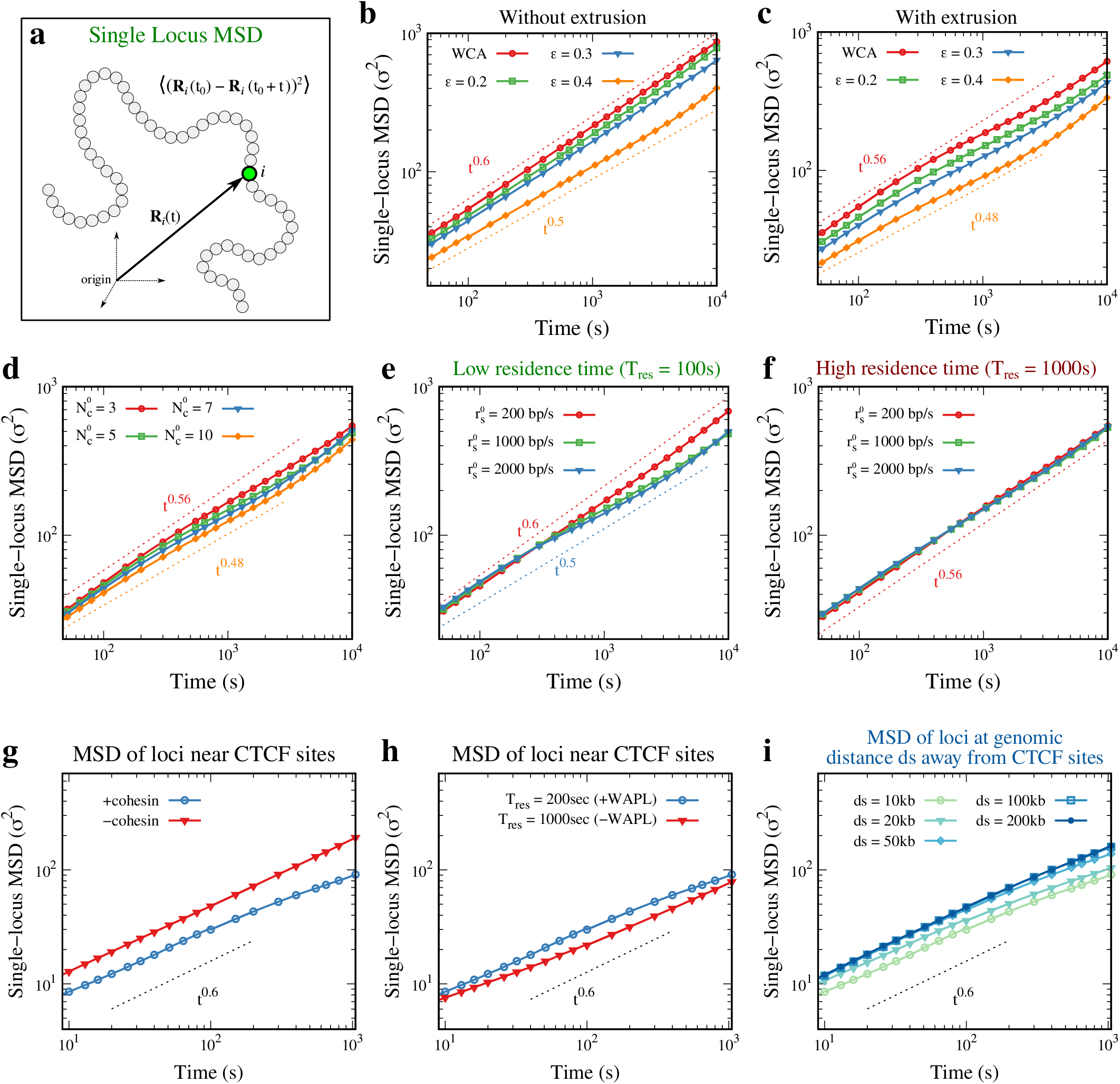
**a** Schematic representing the calculation of single-locus MSD. **b-c** Single-locus MSD as a function of time for different intra-chromatin comparisons is shown for (**b**) a polymer without loop extrusion mechanism and (**c**) polymer with loop extrusion. **d** Single-locus MSD versus time is plotted for different values of average number of cohesins. **e-f** Effect of sliding rate of cohesin on single-locus MSD is shown for (**e**) low residence time and (**f**) high residence time of cohesin complexes. **g** The single-locus MSD of specific loci located near the CTCF sites is plotted in the presence and absence of cohesin. **h** The single-locus MSD of specific loci near the CTCF sites is plotted for low and high values of residence time of cohesin (equivalent to +WAPL and *−* WAPL experimental conditions). **i** The MSD of specific loci genomic distance *ds* away from the CTCF site are plotted as a function of time. In all the plots, apart from the parameters that we vary in each plot, all other parameter values are fixed to an intermediate value: 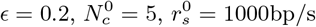 and *T*_res_ = 200s.

First, we check the scaling of MSD with time for polymer without loop extrusion mechanism (Fig. 4b). The polymer with WCA potential shows a scaling of *α* = 0.6 while an attractive LJ polymer with *ϵ* = 0.4 has a scaling of *α* = 0.5. The Rouse dynamics predicts the value of the exponent for a self-avoiding walk (SAW) polymer *α* ≈ 0.55 [85]. This shows that the attractive interactions, even without active loop extrusion, reduce the scaling exponent of MSD with time. As we introduce the active loop extrusion mechanism into the system, this scaling reduces further for all corresponding intra-chromatin interactions (Fig. 4c). In the parameter regime we simulated in this figure (for fixed extrusion parameters), *α* varies in the range of 0.48 to 0.56. Next, we check the effect of other extrusion parameters on the single-locus MSD. We find that the increasing average number of cohesins slightly decreases the scaling of MSD with time (Fig. 4d). However, the exponent *α* remains in a narrow range of 0.48 to 0.54 for the parameters used in Fig. 4d. Next, we compare the effect of sliding rates on single-locus MSD for two extreme values of the residence time of cohesin (Fig. 4e and f). As we saw in the previous section, we expect the time dependence of MSD to be affected by the residence time. For the lower value of the residence time (*T*_res_ = 100s), increasing the sliding rate reduces the scaling exponent *α*, while there is no considerable effect of the sliding rate on single-locus MSD when residence time is higher (*T*_res_ = 1000s). This effect is similar to what we observed for relaxation time (Fig. 3e-f). Note that for different sets of parameter values, the exponent *α* varies around 0.5, which is consistent with what is observed in experiments [30, 89, 90]. As done in recent experiments, we also computed single locus MSD of segments very close to the CTCF sites, ⟨ (**R**_*i*_(*t*_0_ + *t*) *−* **R**_*i*_(*t*_0_))^2^⟩ _*i* ∈ {325,470}_, where beads *i* ∈ {325, 470} are close to the CTCF sites. Our results (see Fig. 4) are comparable to what is observed in experiments[60]. For example, in the absence of extrusion (-cohesin), the MSD curve shifts to a higher value compared to the case with extrusion, +cohesin, keeping a similar power-law exponent (Fig. 4g and Fig. S8). This shift suggests that the generalized diffusion coefficient of loci close to CTCF sites increases in the absence of extrusion and is consistent with what is observed in ref.[60]. This is also counter-intuitive because, naively, active ATP-dependent extrusion is thought to make the chromatin more dynamic as if the effective temperature is high. However, the extrusion activity makes the polymer more ordered by forming loops; polymer bead movement gets restricted due to the loop formation. Hence, the activity decreases the effective diffusivity in a non-trivial way. WAPL is known to help in the unloading of cohesin, and hence experiments lacking WAPL (-WAPL) are equivalent to increasing the residence time (*T*_res_) of cohesin. Figs. 4h and S9 show that an increase in residence time, which is equivalent to WAPL depletion, decreases generalized diffusivity, which is consistent with what is reported in experiments (see ref.[60]). In the absence of CTCF, our simulations show higher diffusivity as expected (Fig. S10a-c). In the absence of CTCF, the polymer beads near the CTCF binding sites may not always get anchored in a looped configuration and would move around even though cohesin is present. This would result in higher diffusivity than polymer with CTCF, as observed. However, surprisingly, experiments do not find much change in diffusivity in the absence of CTCF. In our simulations, this is only possible if the extrusion is very slow with low processivity and the CTCF sites are not getting anchored in loops (Fig. S10a). Future experiments can examine this observation further.

Most existing experiments that study the effect of extrusion on chromatin dynamics tag specific loci close to CTCF binding sites. However, chromatin is a heterogeneous polymer, and the dynamics near CTCF locations can differ greatly from those away from CTCF sites. In our simulations, we computed the dynamics of several loci with genomic distance (*ds*) from the CTCF site. The results (see Fig. 4i) show that single locus MSD and diffusivity vary as a function of separation from the CTCF site. Locus close to the CTCF has a lower diffusivity as the loop anchoring at the CTCF location would restrict the dynamics. This prediction from our simulations can be tested in future experiments. Averaging the dynamics of several loci along the chromatin will have a huge contribution from sites that are away from CTCF, hence reducing the locus-specific effects (see Figs. S8, S9 and S10).

Beyond the dynamics of a single locus, to understand the coupled dynamics of two chromatin loci separated by genomic distance *s*, we compute the two-locus MSD given by *M*_2_ = ⟨ (**R**(*s, t*_0_ + *t*) *−* **R**(*s, t*_0_))^2^⟩. Here **R**(*s, t*) is the vector from any bead *i* to bead *j* separated by distance *s* = |*i − j*| *σ* along the polymer contour (see Fig. 5a). At short time (*t < τ*_*r*_), two loci diffuse independently, while at long time scale the MSD saturates to 2 ⟨*R*^2^(*s*)⟩ for given genomic separation (see Fig. 5b) [30]. At short times, we expect *M*_2_ ∼ *t*^*β*^. The experimentally observed scaling of two-locus MSD is *β* ≈ 0.5 [30]. We study *M*_2_ as a function of time for different strengths of intra-chromatin interactions for chromatin polymer with and without extrusion (Fig. 5c-d). Similar to single-locus MSD, the scaling exponent of two-locus MSD with time decreases with increasing attractive interactions (Fig. 5d). A similar decrease in scaling exponent is also found with an increasing number of cohesins 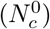 from 3 to 10 (Fig. 5e). Next, we study the effect of the sliding rate on the two-locus MSD for low and high residence time values. For lower residence time, the scaling of two-locus MSD with time decreases with increasing sliding rate, while the sliding rate has minimal effect on the scaling of two-locus MSD when residence time is higher (Fig. 5f-g).

**FIG. 5.**
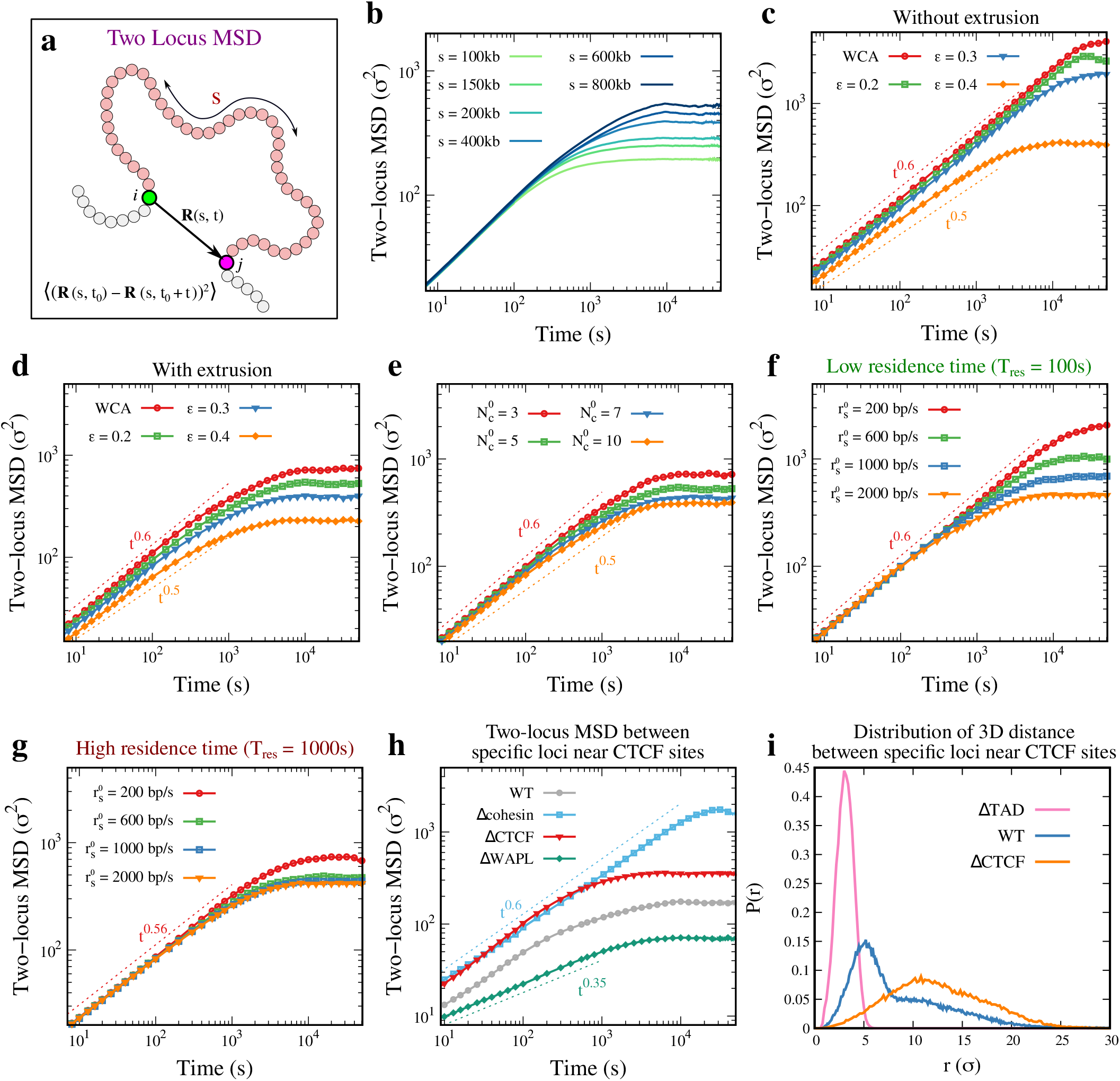
**a** Schematic showing the calculation of two-locus MSD of bead pairs separated by genomic distance *s*. **b** Two-locus MSD averaged over all loci is plotted as a function of time for different values of genomic separation *s*. The MSD values saturate to corresponding ⟨*R*^2^(*s*)⟩ values. **c-d** Two-locus MSD averaged over all loci (*s* = 800 kb) as a function of time for different intra-chromatin interactions is shown for polymers (**c**) without loop extrusion and (**d**) with loop extrusion mechanism. **e** Two-locus MSD averaged over all loci (*s* = 800 kb) versus time is plotted for different values of average number of cohesins. **f-g** Effect of residence time of cohesin on two-locus MSD (*s* = 800 kb) is shown for (**f**) polymer with low processivity and (**g**) high processivity. **h** Two-locus MSD between two specific loci located near CTCF sites is plotted as a function of time for comparison with experimental[59] WT, Δcohesin, ΔCTCF and ΔWAPL conditions. **i** The distribution of 3D distances between two loci near CTCF sites is plotted for comparison with experimental observations[59] of ΔTAD, WT, and ΔCTCF conditions. In all the plots, apart from the parameters that we vary in each plot, all other parameter values are fixed to an intermediate value: 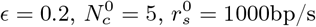 and *T*_res_ = 200s.

We also computed two-locus MSD between two sites that are close to CTCF with a genomic separation of 500kb, ⟨(**R**_72,322_(*t*_0_ + *t*) *−* **R**_72,322_(*t*_0_))^2^⟩, here beads {72, 322} have a separation of 500kb and are close to CTCF sites. We performed separate simulations switching off the effects of CTCF, cohesin, and WAPL independently (Figs. 5h and S11). In the absence of cohesin or CTCF, the diffusivity increases due to the lack of restriction imposed by the loops, similar to the single-locus MSD. In the absence of WAPL, the residence time increases, and the diffusivity of the loci close to CTCF decreases due to the presence of loops. These observations are consistent with what is observed in experiments (see ref.[59]). We also computed the 3D distance distribution between the two loci near CTCF sites (Fig. 5i). Compared to the WT, the peak of the distribution shifts to the right, consistent with an increase in the mean square values in Fig. 5h. If the TAD itself is removed (ΔTAD), the two marked beads near CTCF will have very little mean distance, and the distribution will peak at a smaller value of *r*. All of these are consistent with the experimental observations by Gabriele et al.[59]. We emphasize that all of these results can be different if one examines a location far away from CTCF and is reflected when we present results averaged over locations (see Fig. S11d-f).

## III. CONCLUSIONS

In this work, we used a chromatin model with intra-chromatin interactions and active loop extrusion mechanism to systematically probe the effect of various model parameters on the structure and dynamics of the chromatin. Our model predictions of contact map patterns, 3D distance, and boundary probability are consistent with the experimental observations. We showed how intra-chromatin interactions and cohesin density provide two different mechanisms of chromatin compaction. We demonstrate, contrary to the current notion, that the processivity of the cohesin extruders is not enough to explain the chromatin structure [33]. Beyond the processivity, the compaction and dynamics of the chromatin domains also depend on the interplay between the residence time of cohesin and the polymer segment’s relaxation time. This observation has implications beyond the TADs, as the relative timescales of extrusion activity and polymer relaxation time can affect the complex higher-order structures.

We also investigated how the extrusion activity affects the chromatin dynamics. We observed that the scaling exponent of relaxation time with genomic separation decreases with an increase in the overall extrusion activity and strength of attractive intra-chromatin interactions. The increased extrusion activity can be through higher processivity or cohesin density. However, if the residence time is long enough, such that the cohesins form the complete loop and remain there, then the increase in processivity has no effect on the relaxation time. A similar effect of the extrusion activity was observed on the scaling of single-locus and two-locus MSD with time.

This work provides several testable predictions. We predict how cohesin binding/dissociation rate affects static as well as temporal quantities (see Fig.2, 3g-h, 4e-f, and 5f-g). This can be tested in experiments with WAPL, NIPBL, and PDS5 mutants. Our simulations predict very different outcomes when the extrusion time scale is slower or faster than the relaxation time (Fig. 2e). The chromatin polymer is heterogeneous, and we predict how dynamic behavior of various mutants depends on the precise genomic location where one measures it (dynamics near CTCF sites versus dynamics of regions away from the domain boundary) as shown in Fig. 4i. Another important testable prediction is that dynamics of chromatin in *Drosophila* would be different in the presence or absence of cohesin (see Fig. 3i). We predict a similar phenomenon is also possible in some specific regions of mammalian chromosome, where intra-domain interactions compete with loop extrusion. All of these also have implications for gene regulation.

One of the limitations of our work is that we do not explicitly include dynamic properties of the medium in which the polymer is embedded. Since chromatin *in vivo* is in a highly crowded environment, the hydrodynamic interactions are expected to be screened, and a study without hydrodynamics may be reasonable. However, we were able to establish how extrusion parameters contribute to the trends of increase or decrease in the scaling exponents of dynamic quantities. Our comprehensive analysis provides important insights into how various parameters of the extrusion mechanism can affect the structure and dynamics of chromatin loci.

## IV. METHODS

We model chromatin as a coarse-grained bead-spring polymer consisting of *N* beads connected by harmonic springs. The whole polymer represents the chromatin region of size 1 Mb, with each bead of size 2 kb. At this resolution, chromatin can be regarded as a flexible polymer [14, 21]. We integrate the loop extrusion process into our 3D polymer model by introducing rate-dependent binding, dissociation, and sliding of cohesins on the chromatin polymer (Fig. 1a). These rate processes update the positions of bound cohesins, which in turn affect the 3D polymer configurations. The total energy of the polymer is given by

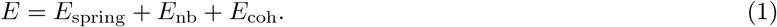

The first term *E*_spring_ in the above equation represents the spring energy between the neighboring beads along the polymer, the second term *E*_nb_ represents the non-bonded interactions between chromatin beads, and the third term *E*_coh_ captures the energy due to the cohesin complexes connecting monomer beads at their anchor positions during the extrusion process (see detailed implementation of loop extrusion mechanism below). The spring energy *E*_spring_ is given by

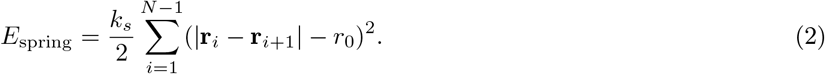

Here, **r**_*i*_ is the position vector of *i*^*th*^ bead, *r*_0_ is the equilibrium bond length, and *k*_*s*_ is the spring constant.

We have two types of models depending on the nature of the non-bonded interactions between the chromatin beads: 1. All-bead uniform interaction model and 2. Intra-domain bead attraction model.

### All-bead uniform interaction model

In this model, all the non-bonded beads interact with each other through same interactions (*E*_nb_). We model this using the attractive Lennard-Jones (LJ) potential or the repulsive Weeks-Chandlers-Andesron (WCA) potential [74]. The LJ potential has a repulsive core and short-range attractive interaction, which is given by

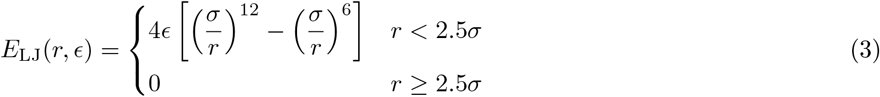

Here, *σ* is the diameter of the monomer bead, while the LJ energy parameter *ϵ* denotes the depth of the potential and controls the strength of attractive interactions. The WCA potential consists of only the repulsive part of LJ potential [74] and is given by

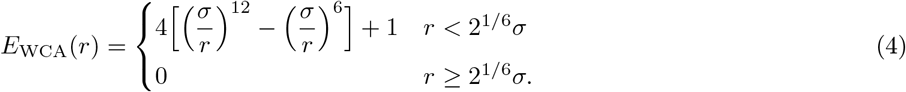

### Intra-domain bead attraction model

In this model, the non-bonded interactions (*E*_nb_) between the chromatin beads that are within the same domain are modeled through the attractive LJ interaction (*E*_LJ_(*r, ϵ*_intra_)), while the inter-domain bead interactions are modeled using the WCA potential (*E*_WCA_(*r*)). The domains are considered based on the CTCF locations, which mark the domain boundaries.

### Loop extrusion mechanism

We have modeled the loop extrusion mechanism using rate events, where cohesin can bind, dissociate, and slide along the chromatin (Fig. 1a). We also consider four CTCF sites bound at the following genomic locations: 39.216 Mb, 39.35 Mb, 39.84 Mb, and 40.146 Mb. We have chosen these locations by comparing the TAD boundary locations from the Hi-C data[91] and the CTCF ChIP-seq data[92] (see Fig. S1). Since there are multiple CTCF motifs near the TAD boundaries, we assume that the CTCF is always bound at these locations. These sites are considered permanently bound by CTCF unless specified otherwise. The bound cohesin occupies two beads on the chromatin polymer, representing the two anchor sites. The effect of cohesins is purely through the harmonic spring contact between the two chromatin beads occupied by the cohesin anchor sites (last term in Eq.1). When a cohesin binding event occurs, a binding site is chosen randomly so that the two adjacent beads are not occupied by either CTCF or other cohesins. Each anchor site of bound cohesins independently slides in opposite directions to form a loop. If one of the anchor sites of cohesin is blocked by a CTCF barrier or another cohesin, then that anchor site does not slide, while the opposite anchor site can keep moving. Since each cohesin occupies two beads on the chromatin polymer, the number of occupied sites by *N*_*c*_ number of cohesins is 2*N*_*c*_. If there are 2*N*_*c*_ number of cohesin bound sites and *N*_*f*_ number of free binding sites available at a given moment, then the binding and dissociation rates of cohesin are given by the following equations:

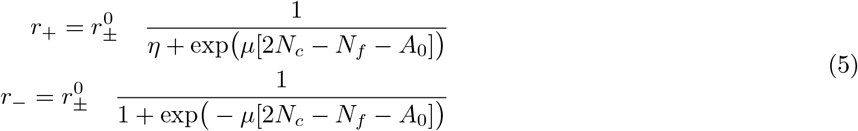

Here, 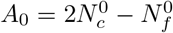 is an asymmetry parameter where 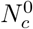 and 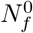 are the average number of bound cohesins and average number of free sites, respectively. The value of the parameter *η* is chosen so as to satisfy the detailed balance at steady state. Therefore, the condition at steady state 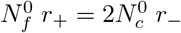 gives us the value 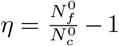. The parameter *µ* controls the fluctuations around the steady state value, and 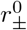 is the attempt rate for the binding/dissociation process. When a binding event is chosen, cohesin binds at a randomly chosen available site occupying the adjacent beads (*i* and *i* + 1). The cohesin complexes slide along the chromatin polymer at a rate 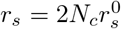, such that both the anchor sites move in opposite directions provided that the site is not occupied by any other cohesin or CTCF. 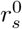 is the intrinsic rate of sliding of each cohesin anchor. The two anchor sites of each cohesin are connected by a harmonic spring represented by the last term in Eq.1, which is given by

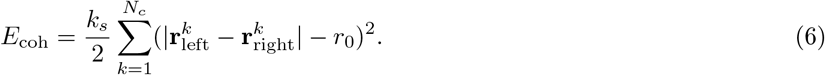

Here 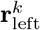 and 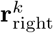 are the position vectors of beads where the corresponding left and right anchor sites of the *k*^*th*^ cohesin are present.

We simulate the system using the Monte-Carlo algorithm [73]. We have three types of moves: 1. change the position of a randomly selected bead, 2. perform binding and dissociation events of cohesin complex with rate *r*_*±*_, and 3. slide each cohesin anchor with rate *r*_*s*_. The position move is performed by adding uniform random numbers *r*_*n*_ ∈ [ *−δr*, +*δr*] to the x, y, and z coordinates of the chosen bead. The position move is accepted based on the Metropolis criterion with the total energy computed using Eq. 1. A single Monte-Carlo step (MCS) consists of *N* such moves. The binding and dissociation moves are attempted *N* times during a single MCS. From the instantaneous number of bound cohesin, we get the values of rates *r*_*±*_ from Eq. 5. Similarly, the sliding event is attempted *N*_*c*_ number of times (for both left and right anchor sites) with rate 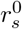 during a single MCS (resulting in an effective sliding rate 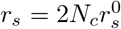). The corresponding probability of accepting these events is given by *P*_*±*_ = min(1, r_*±*_) and *P*_*s*_ = min(1, r_s_).

In our Monte Carlo simulations, we have ensured that all the moves are simple physical diffusive moves, i.e., all conformational changes are based on moves at single bead level. In an MCS an attempt is made to move every bead by a distance *δr* in three dimensions. We do not use any multi-bead level moves like segmental rotations or other unphysical moves. Hence, the dynamics that we compute from purely diffusive moves are indeed correct and can be mapped to real-time[93]. We have verified this by computing the dynamic quantities for a Rouse chain and comparing it with the known analytical results (Fig. S12).

### Initialization and sampling

We initialize the polymer trajectories in self-avoiding random walk configurations. We simulate the system for 10^8^ MCS and sample configurations every 50,000 MCS. All the steady state properties are computed by excluding the data for initial 2×10^7^ MCS. For each parameter set, we have simulated 15 such trajectories.

### Parameters

All simulations are performed using dimensionless units, with length measured in bead diameters (*σ*), energy in units of *k*_*B*_*T*, and time in units of MCS. The mapping of these dimensionless units to real units is discussed in the Supplementary Note A. The polymer size is *N* = 500. In Eq. 2 and Eq. 6, the spring constant is set to *k*_*s*_ = 100 and the equilibrium distance to *r*_0_ = *σ*. We vary the strength of attractive interactions in Eq. 3 within the range *ϵ* ∈ [0.2, 0.4]. The parameter controlling the fluctuations of the number of bound cohesins in Eq. 5 is *µ* = 10. The mean number of cohesins is varied in the range 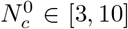. The attempt rate for the binding/dissociation process in Eq. 5 varies within 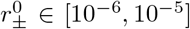, while the sliding rate varies within 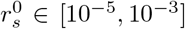. For position moves in Monte Carlo algorithm, *δr* is set to 0.5*σ*. The conversion of these dimensionless quantities into real units for comparison with experiments is discussed in Supplementary Note A.

## V. DATA AVAILABILITY

Published Hi-C data[91] and CTCF ChIP-seq data[92] used in this study are available in the Gene Expression Omnibus (GEO) database with accession numbers GSE130275 and GSE96107. Relevant data generated from this study are included in this article’s Figures, text, and supplementary information.

## VI. ACKNOWLEDGMENTS

We acknowledge the National Supercomputing Mission (NSM) for providing computing resources of ‘PARAM Brahma’ at IISER Pune, which is implemented by C-DAC and supported by the Ministry of Electronics and Information Technology (MeitY) and Department of Science and Technology (DST), Government of India. This work was benefitted from discussions at the International Centre for Theoretical Sciences (ICTS) via the program Interdisciplinary aspects of chromatin organization and gene regulation (code: ICTS/chromatin2024/09). S.K. acknowledges fellowship support from the CSIR, India.

## SUPPLEMENTARY INFORMATION

### S1. Supplementary Notes

#### A Conversion of simulation parameters to real units for comparison with biological systems

The simulation time is denoted in Monte Carlo steps (MCS). To relate simulation time to real-time, we compare the relaxation time of a chromatin segment of size 1000kb from our simulation (with intermediate values of extrusion parameters) to the experimentally observed value from Bruckner et al. [1]. The relaxation time of the 1000kb segment from simulations is *τ*_*r*_ ≈ 10^6^ MCS, while experiments show *τ*_*r*_ ≈ 1000 s, indicating 1 MCS ≈ 1 ms. We utilize this conversion to estimate the values of extrusion parameters in real units below.

- **Extrusion speed of cohesin:** The sliding rate of individual anchors of cohesin 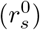 in simulations ranges from 10^*−*5^ beads/MCS to 10^*−*3^ beads/MCS, where one bead corresponds to a chromatin segment of size 2kb. Converting this to real units yields extrusion speeds (twice the speed of extrusion of individual cohesin anchors) ranging from 40 bp/s to 4 kb/s. This covers the experimentally observed range of extrusion speeds from 100 bp/s to 2 kb/s [2–5].
- **Residence time of cohesin:** The residence time of cohesin 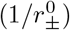 in simulations ranges from 10^5^ MCS to 10^6^ MCS. In real units, this corresponds to a range of 1 minute to 17^*±*^ minutes, consistent with the experimentally observed residence time ranging from 5 minutes to 20 minutes [6–8].
- **Cohesin density:** In simulations, cohesin density ranges from 3 per Mb to 10 per Mb. This encompasses the experimentally measured density of 1 per 240 kb (approximately 4 per Mb) [4].
- **Processivity of cohesin:** The processivity of an individual anchor of cohesin in the simulation is defined as 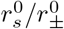, which varies from 10 beads to 100 beads (up to 1000 beads, only for 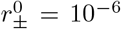. Therefore, the processivity of cohesin (twice that of the individual anchor) in real units ranges from 40 kb to 400 kb. This range includes the experimentally observed processivity value of 150 kb [4].

#### B Additional details on quantities measured

- **Contact probability:** The contact probability (*P*_*ij*_) quantifies the likelihood of interaction between two beads, denoted by indices *i* and *j*. It is determined as the average occurrence of contact overall polymer configurations, computed using the following formula:

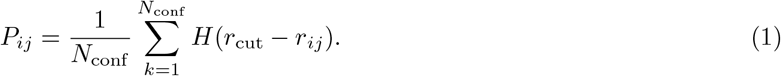

Here, *N*_conf_ represents the total number of polymer configurations, *H*(*r*) denotes the Heaviside step function,*r*_*ij*_ is the distance between the beads *i* and *j*, and *r*_cut_ signifies the cutoff distance, below which two beads are considered in contact. We use *r*_cut_ = 1.5*σ* as the cutoff for defining contact.

- **Boundary probability:** It is the probability of finding the boundary of a locally collapsed TAD-like domain, at a given genomic location. We have adapted an algorithm from Bintu et al. to calculate this probability based on an ensemble of configurations obtained from our simulations [9]. To determine the boundary probability, we first identify boundary locations by analyzing individual chromatin conformations and then average over all configurations.

To locate the boundaries within individual configurations, we initially create a distance map by computing the 3D distance *r*_*ij*_ between all pairs of beads *i* and *j*. We then identify the beginning and end of a domain boundary separately. To determine the start of a domain, we compare the average 3D distances in a left stripe (*S*_*l*_(*k*)) and a right stripe (*S*_*r*_(*k*)) with respect to the genomic coordinate *k* (see Fig. S13). Mathematically, this is expressed as:

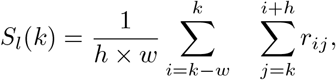

and

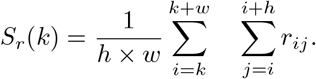

Here, *h* and *w* are the height and width of the stripes, respectively. At the start of a domain, the average 3D distance in the left stripe (*S*_*l*_) will be higher than that in the right side (*S*_*r*_). The genomic coordinates where this ratio *S*_*l*_*/S*_*r*_ has a local maximum represent the boundaries at the start of domains in a given polymer conformation.

Similarly, the genomic coordinate *k* where the ratio of average 3D distances in the top stripe (*S*_*t*_(*k*)) to the bottom stripe (*S*_*b*_(*k*)) is maximum signifies the end of a domain. The average 3D distance in these stripes is calculated as:

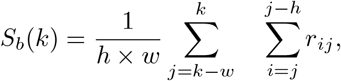

and

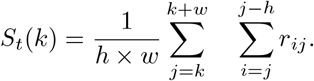

Note that, depending on the number of domains, there can be multiple boundaries in a single polymer conformation. We perform this analysis over an ensemble of polymer conformations to get the boundary probability as a function of genomic location. The values of the parameters *h* and *w* are chosen empirically as *h* = 200kb and *w* = 40kb. The exact values of the boundary probabilities may vary depending on these values; however, the qualitative features remain the same.

- **Radius of gyration:** The radius of gyration *R*_*g*_ is used to quantify the size of the chromatin polymer. The average radius of gyration of the polymer is given by

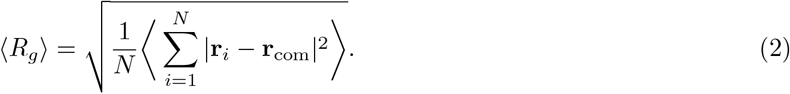

Here, the angled brackets denote the time and ensemble average over steady state configurations, **r**_*i*_ is the position vector of *i*^*th*^ bead, and **r**_*com*_ denotes the position vector of the center of mass of the polymer, which is calculated as:

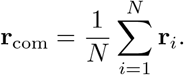

### S2. Supplementary Figures

**FIG. S1.**
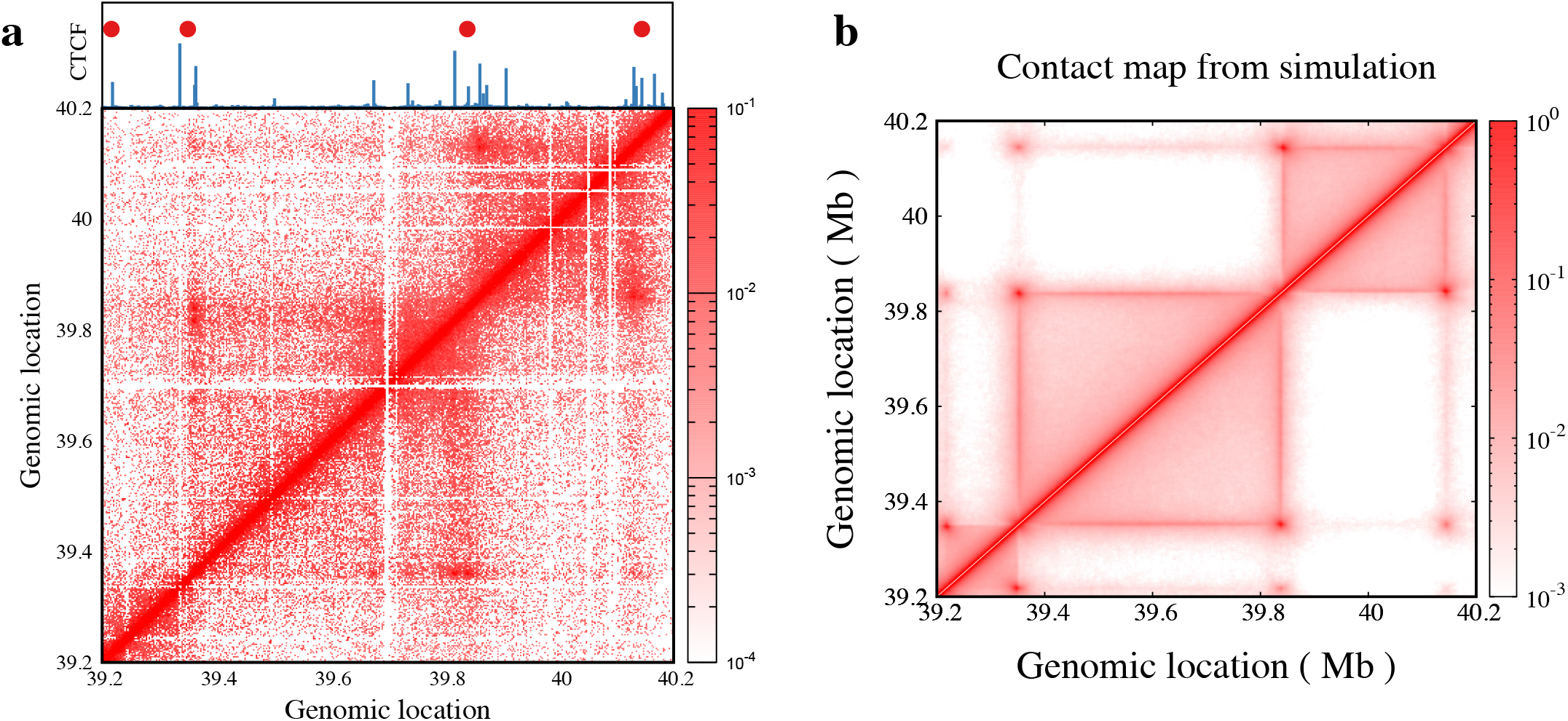
**a** Experimental ontact map data for WT mESC cells[10] is plotted for the genomic region chromosome 15: 39.2 Mb to 40.2 Mb as a heatmap. The ChIP-seq data of CTCF binding[11] for the corresponding region is shown on the top (blue line). The red filled circles represent the permanent CTCF binding sites chosen in the simulation marking the boundary locations of TADs.**b** Contact map from simulation for all-bead uniform interaction model with extrusion is plotted for intermediate extrusion parameters: 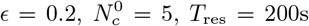 and 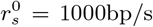. The minimal model captures the TAD structures observed in experiments.

**FIG. S2.**
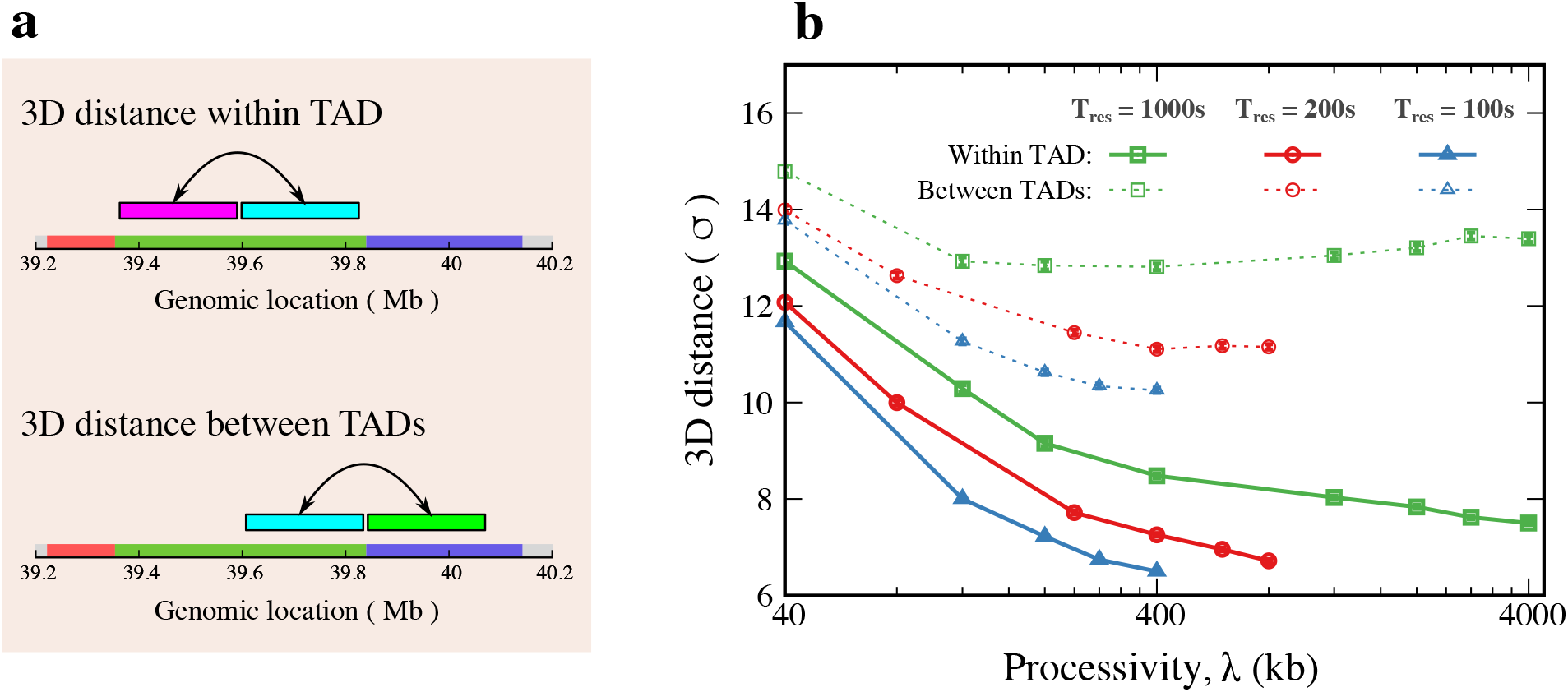
**a** The schematic shows the chromatin segments that lie within same TAD or different TADs for the measurement of 3D distance. **b** Average 3D distance between center of mass of segments that lie within same TAD and between different TADs, as shown in (**a**) is plotted as a function of *λ* for different values of *T*_res_. The values of remaining parameters are fixed (*ϵ* = 0.2 and 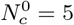) for all the plots.

**FIG. S3.**
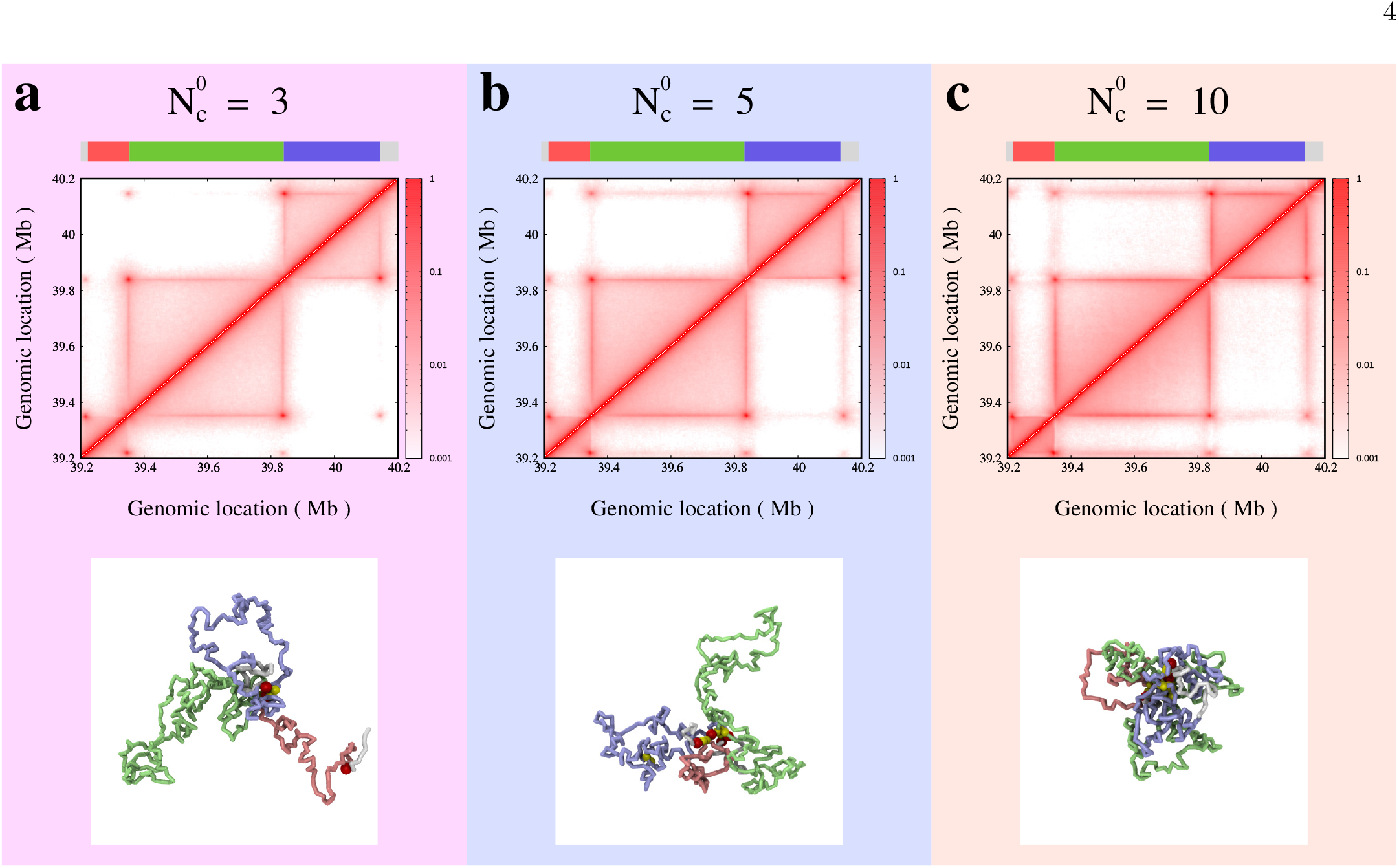
**a-c** Contact maps of polymer with loop extrusion mechanism for different mean number of extruders (**a**) 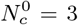, (**b**) 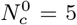, and (**c**) 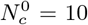 are shown in top row. The bottom row shows representative snapshots for extruder densities. The polymer segments are colored to distinguish different TADs as shown in the color strip on top. The other parameter values used are *ϵ* = 0.2, *T*_res_ = 200s and 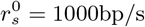.

**FIG. S4.**
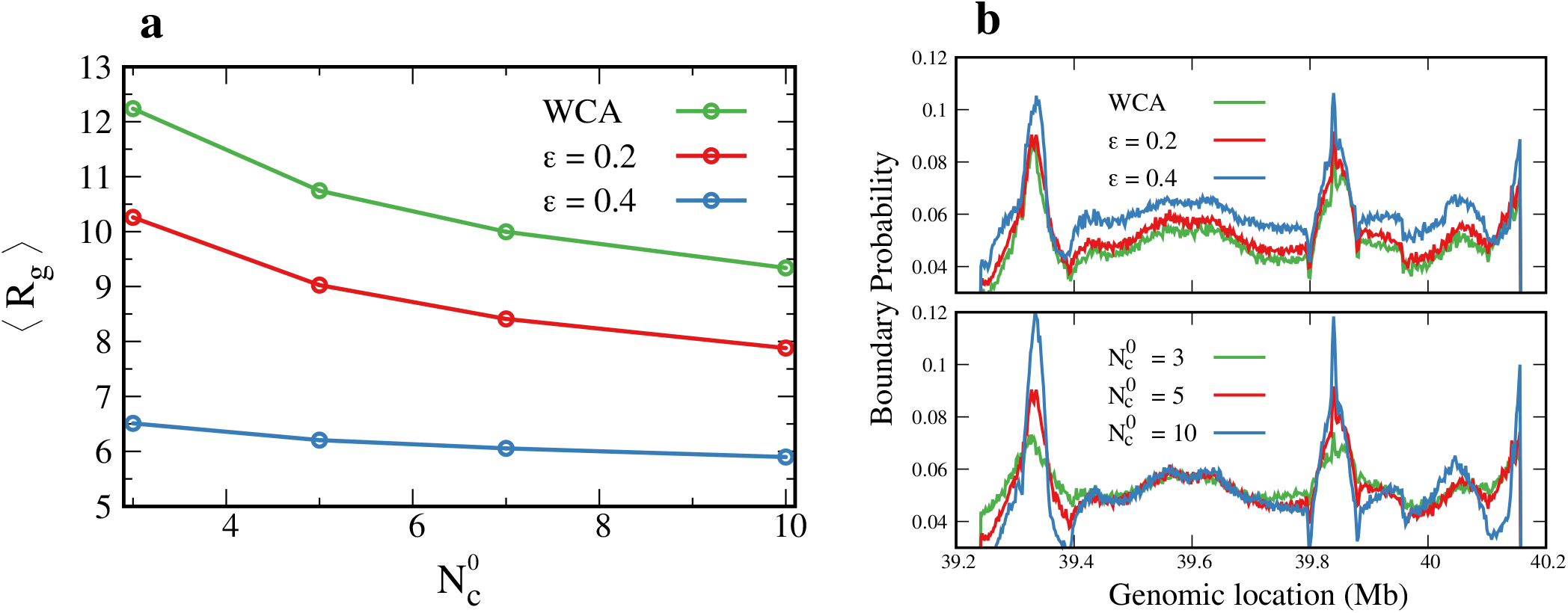
**a** The average radius of gyration is plotted as a function of average number of extruders for different intra-chromatin interactions. **b** The boundary probability as a function of genomic location is shown for different intra-chromatin interactions (top) and for different average number of extruders (bottom). The values of parameters not mentioned in plots are fixed to intermediate values: 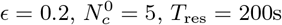 and 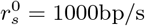.

**FIG. S5.**
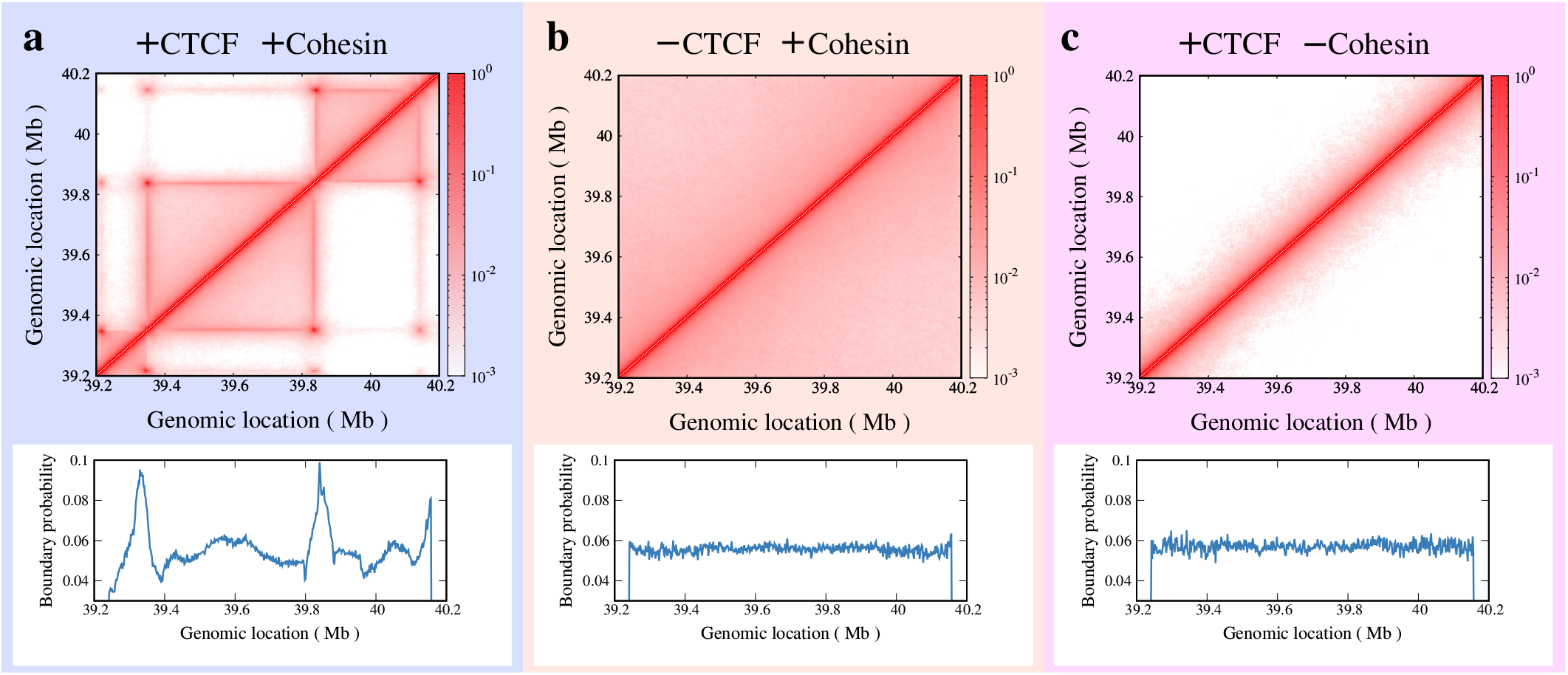
**a-c** Comparison of contact map and boundary probability plots for chromatin polymer with (**a**) both CTCF and cohesin extruders present, (**b**) CTCF absent but cohesin present and (**c**) CTCF present but cohesin absent (i.e. no extrusion). The parameter values used are 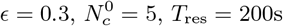 and 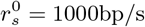.

**FIG. S6.**
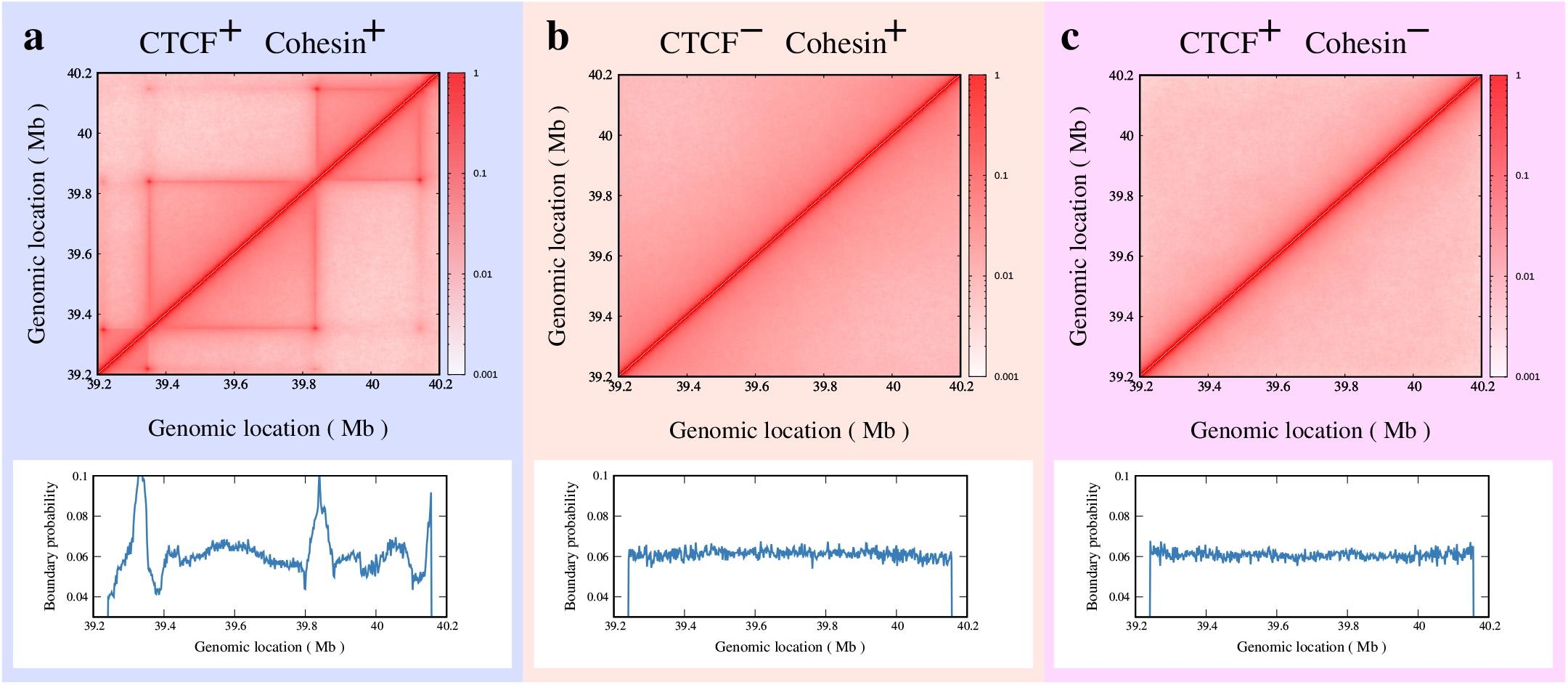
**a-c** Comparison of contact map and boundary probability plots for chromatin polymer with (**a**) both CTCF and cohesin extruders present, (**b**) CTCF absent but cohesin present and (**c**) CTCF present but cohesin absent (i.e., no extrusion). The parameter values used are 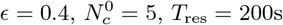 and 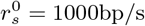.

**FIG. S7.**
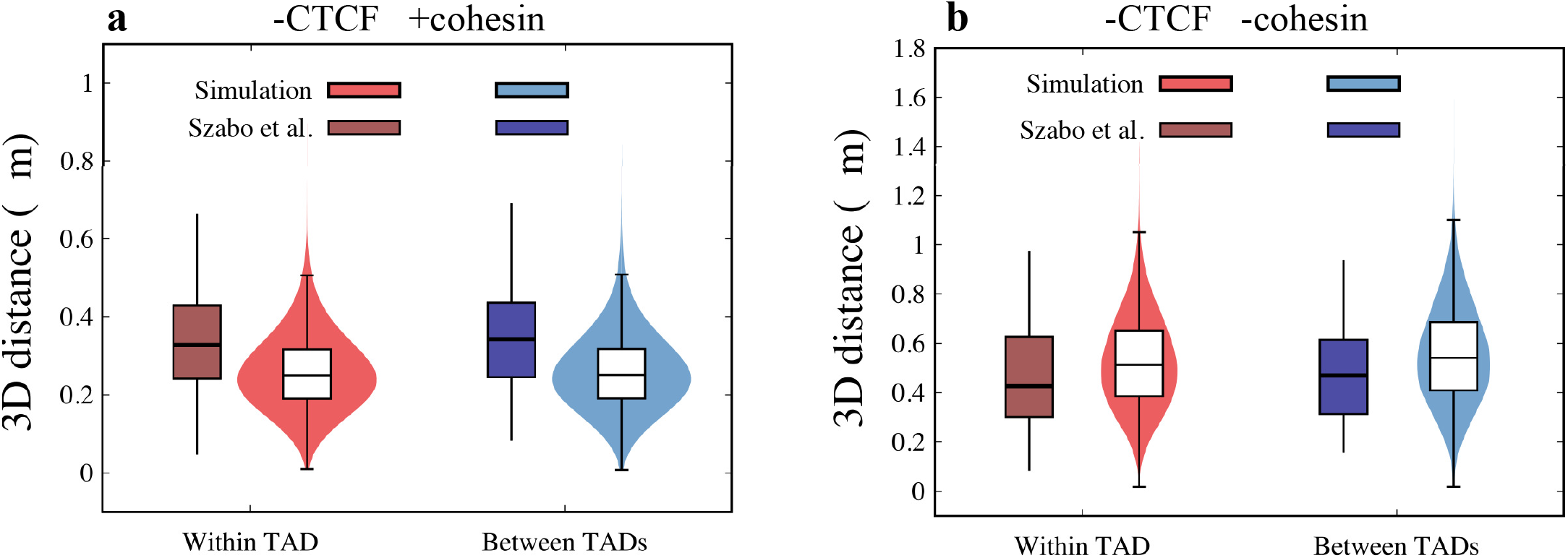
**a-b** The box-plot compares the 3D distance between two segments that are within the same TAD or between the neighboring TADs for (**a**) *−*CTCF +cohesin and (**b**) *−*CTCF *−*cohesin conditions. The distribution of the data points from simulation (*σ* = 40nm) is plotted as a violin-plot. The 3D distance observed from our simulations is comparable to the experiments by Szabo et al. [12]. The parameter values used are 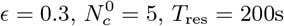 and 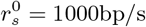.

**FIG. S8.**
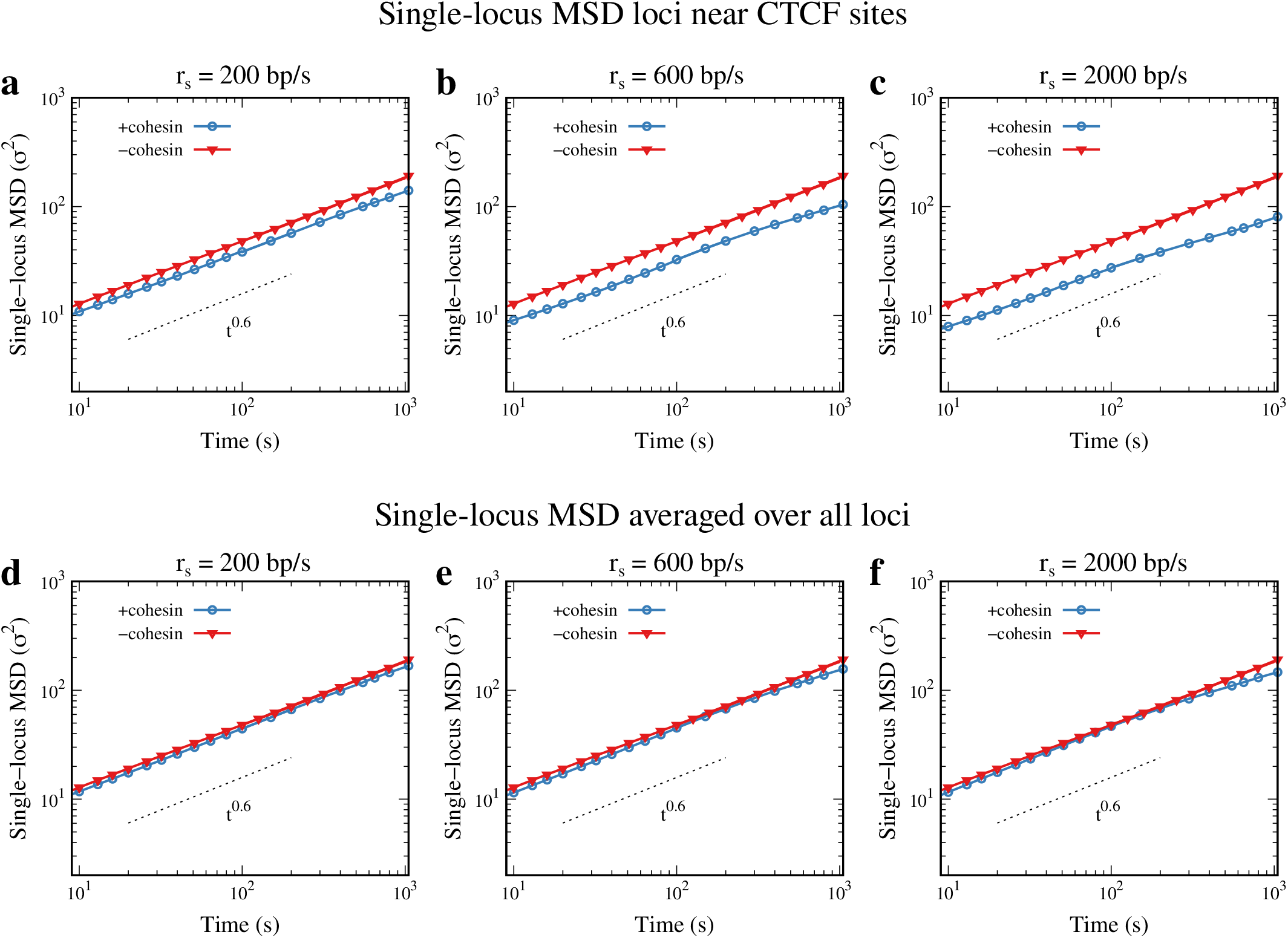
**a-c** Single-locus MSD of loci close to the CTCF sites is compared in the presence and absence of cohesin for different sliding rates. The difference in the single-locus MSD is minimal for smaller sliding rate (**a**) *r*_*s*_ = 200bp/s. With increasing sliding rate (**b** *r*_*s*_ = 600bp/s and **c** *r*_*s*_ = 2000bp/s), the MSD curve in the presence of cohesin shifts downward. **d-f** Single-locus MSD averaged over all loci is compared in the presence and absence of cohesin for different sliding rates. The change in the single-loucs MSD values when averaged over all loci is very minimal for all sliding rates as compared to the single-locus MSD of loci near CTCF sites.

**FIG. S9.**
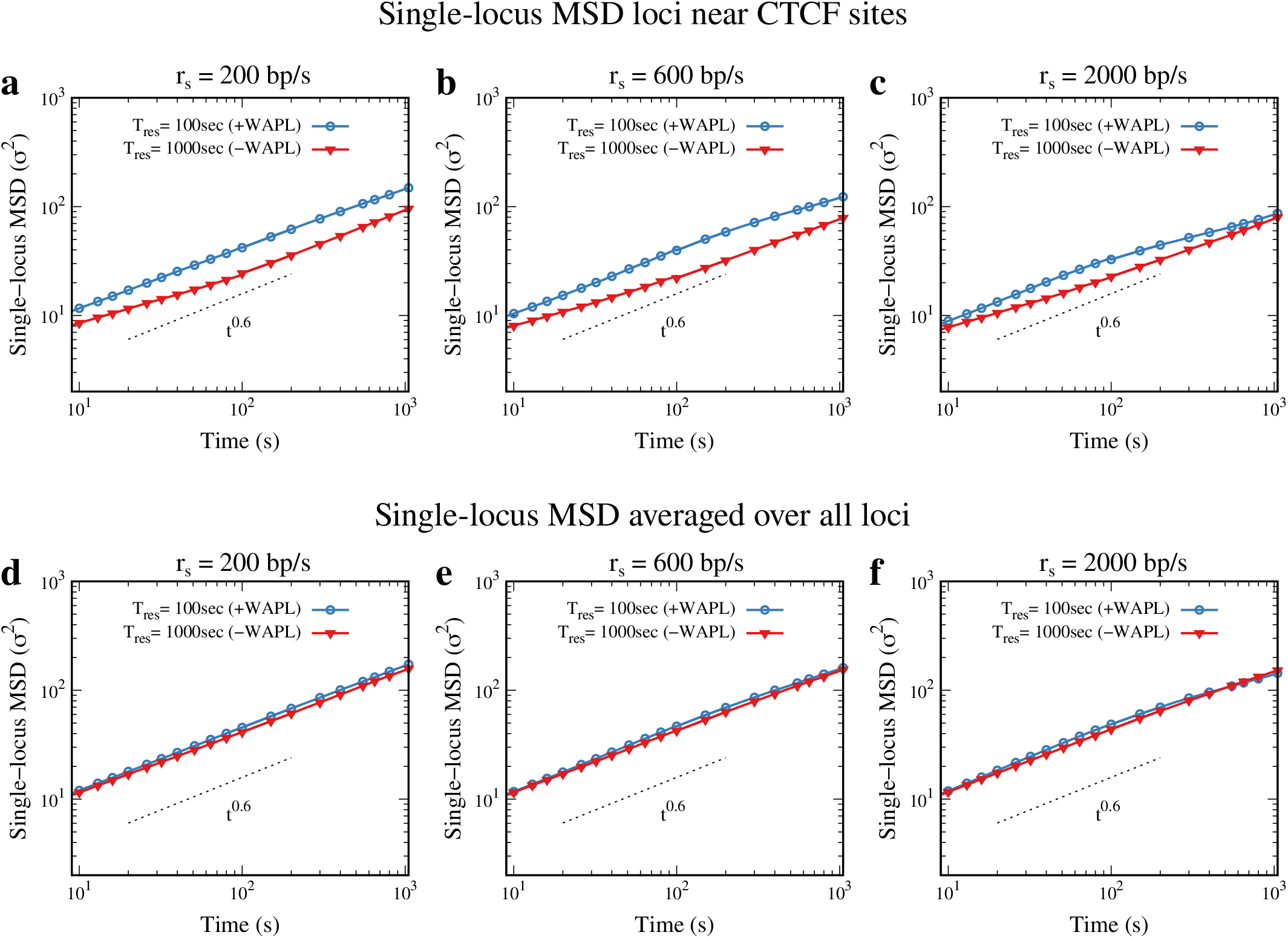
**a-c** Single-locus MSD of loci close to the CTCF sites is compared in the presence and absence of WAPL (different values of residence time *T*_res_ of cohesin) for different sliding rates. The difference in the single-locus MSD is higher for smaller sliding rate (**a**) *r*_*s*_ = 200bp/s. With increasing sliding rate (**b** *r*_*s*_ = 600bp/s and **c** *r*_*s*_ = 2000bp/s), the difference in the MSD curves in the presence and absence of WAPL decreases. **d-f** Single-locus MSD averaged over all loci is compared in the presence and absence of WAPL for different sliding rates. The change in the single-loucs MSD values when averaged over all loci is very minimal for all sliding rates as compared to the single-locus MSD of loci near CTCF sites.

**FIG. S10.**
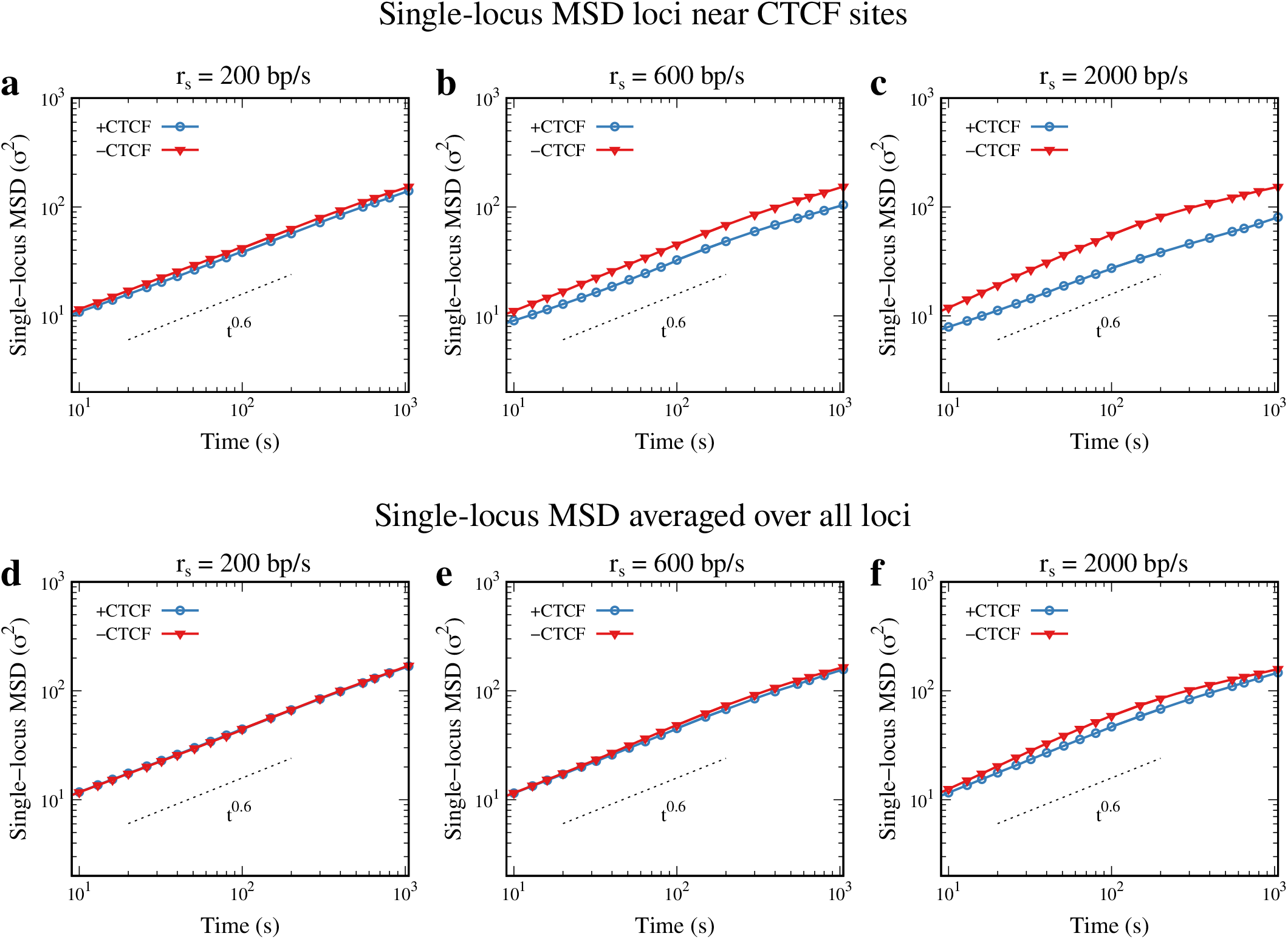
**a-c** Single-locus MSD of loci close to the CTCF sites is compared in the presence and absence of CTCF complex for different sliding rates. The difference in the single-locus MSD is minimal for smaller sliding rate (**a**) *r*_*s*_ = 200bp/s. With increasing sliding rate (**b** *r*_*s*_ = 600bp/s and **c** *r*_*s*_ = 2000bp/s), the MSD curve in the presence of CTCF shifts downward. **d-f** Single-locus MSD averaged over all loci is compared in the presence and absence of CTCF complex for different sliding rates. The change in the single-loucs MSD values when averaged over all loci is very minimal for all sliding rates as compared to the single-locus MSD of loci near CTCF sites.

**FIG. S11.**
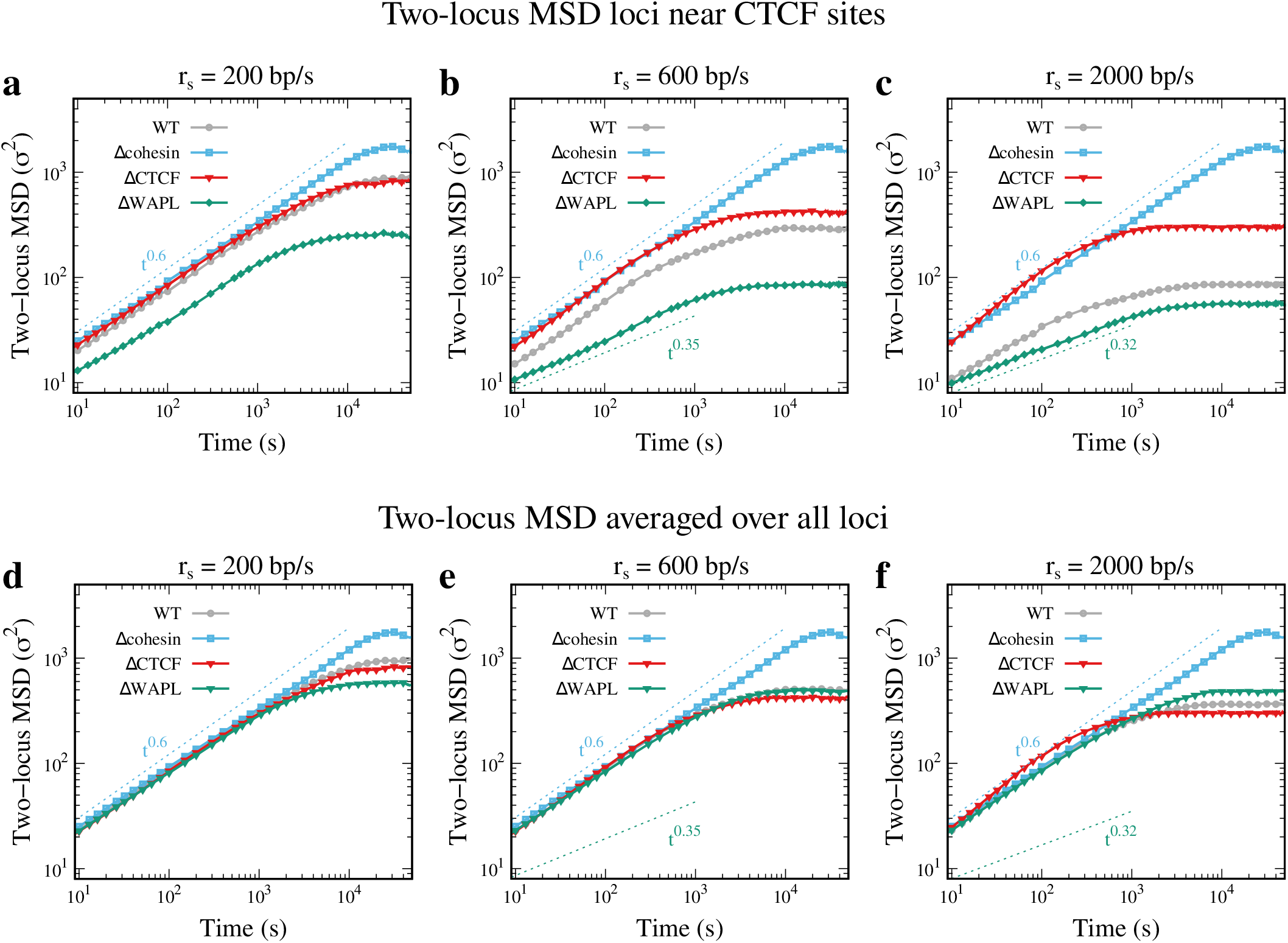
**a-c** Two-locus MSD of loci near the CTCF sites is compared between WT (+CTCF +cohesin), Δcohesin, ΔCTCF and ΔWAPL (higher residence time of cohesin) conditions, with increasing sliding rate *r*_*s*_. The two-locus MSD for Δcohesin is the highest followed by ΔCTCF, WT and ΔWAPL, with the difference increasing with increasing sliding rate *r*_*s*_. **d-f** Two-locus MSD averaged over all loci is compared between WT (+CTCF +cohesin), Δcohesin, ΔCTCF and ΔWAPL (higher residence time of cohesin) conditions, with increasing sliding rate *r*_*s*_. The change in the two-loucs MSD values when averaged over all loci is very minimal for all sliding rates as compared to the two-locus MSD of loci near CTCF sites.

**FIG. S12.**
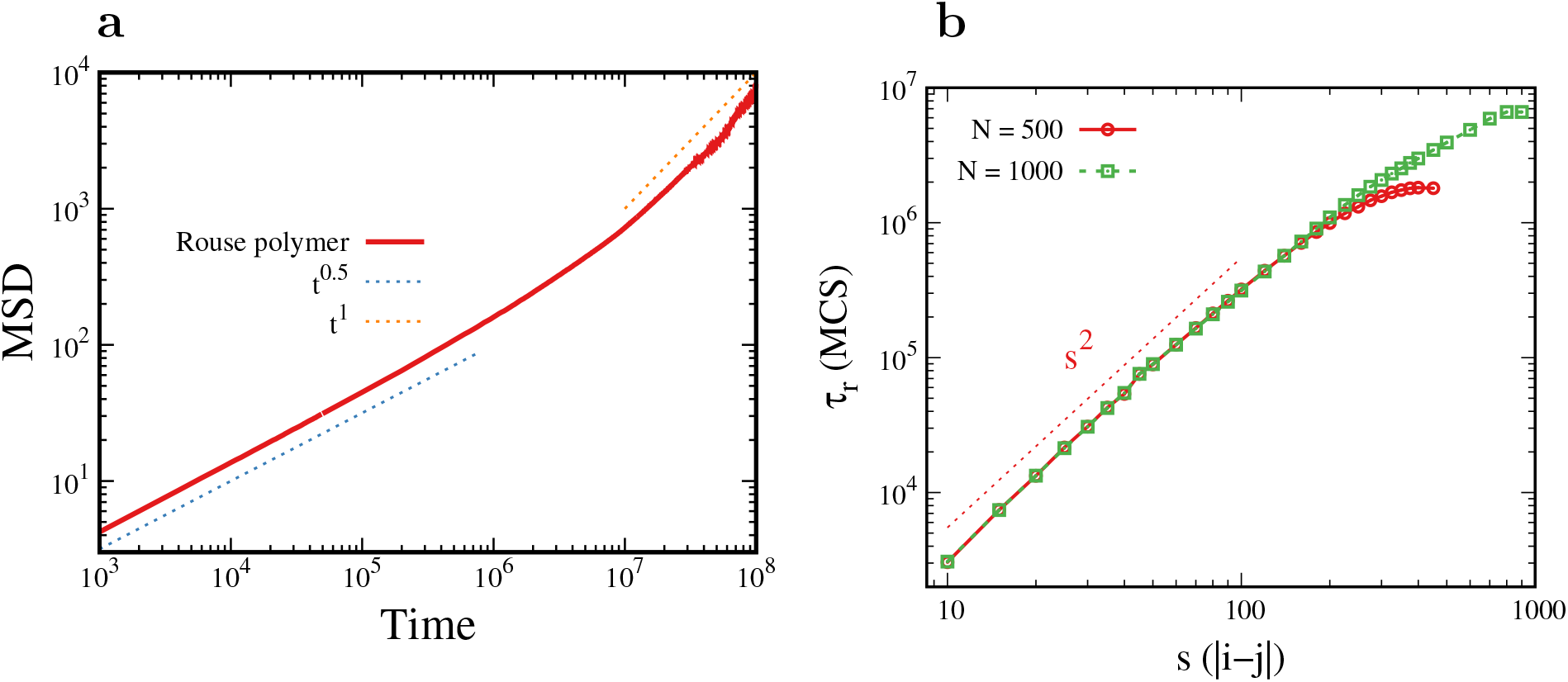
**a** Mean-square displacement of monomer in rouse polymer is plotted as a function of time (MCS). The scaling is consistent with the known analytical results. **b** The relaxation time (*τ*_*r*_) as a function of contour separation is plotted for Rouse polymers with *N* = 500 and *N* = 1000. The scaling is consistent with the known analytical results for Rouse polymer. The slight decrease observed towards the end, where contour separation is close to polymer size, decreases slightly due to the finite-size effects.

**FIG. S13.**
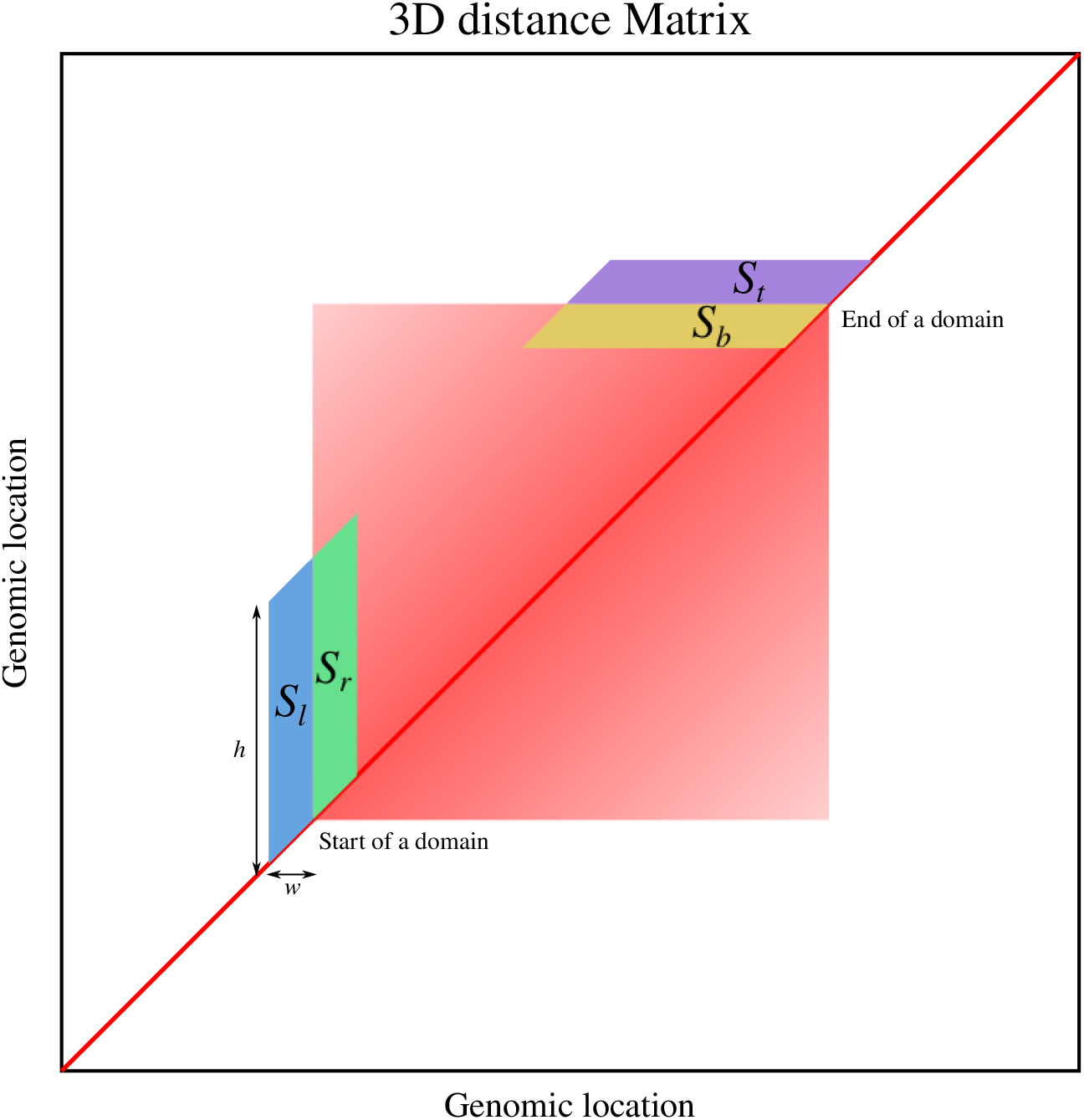
The schematic representation of boundary analysis. The 3D distance matrix from individual configurations is used to identify the locations across which there is maximal change in the average 3D distance. The stripes of size *h* × *w* are used to compute average 3D distance across the genomic coordinates. The ratio of average 3D distances from left stripe to right stripe (*S*_*l*_*/S*_*r*_) shows local maxima at the genomic coordinate where start of a domain. Similarly, the ratio *S*_*t*_*/S*_*b*_ has a local maxima at genomic coordinate where a domain ends.

### Supplementary movie caption

Movie S1. The movie compares the steady state trajectories of polymers exhibiting high processivity 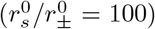 under two conditions: one with a low residence time 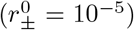 of cohesins on the left, and the other with a high residence time 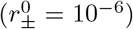. The configurations are shifted to the center of mass frame of reference for visualization purposes. Other parameter values are *ϵ* = 0.2 and 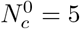.

